# Carbon monoxide oxidation expands the known metabolic capacity in anaerobic methanotrophic consortia

**DOI:** 10.1101/2025.09.21.677609

**Authors:** Yongzhao Guo, Daniel R. Utter, Ranjani Murali, Victoria J. Orphan

## Abstract

Efficient energy metabolism is essential for microorganisms living on the thermodynamic edge. Consortia of anaerobic methane-oxidizing archaea (ANME-2) and sulphate-reducing bacteria (SRB) represent globally relevant syntrophic associations capable of growing with minimal amounts of free energy and can persist when methane becomes limiting. Their potential for physiological plasticity, including the use of electron donors beyond methane is poorly understood. Carbon monoxide (CO) has been reported in seep environments and represents a thermodynamically favourable alternative electron donor due to its low reduction potential. We demonstrate that environmental ANME-SRB consortia can oxidize CO in the absence of methane, in anoxic microcosm experiments using a combination of stable isotope geochemical tracers, metatranscriptomics, and single cell activity measurements (FISH–nanoSIMS). The oxidation of CO was coupled with sulphate-reduction by syntrophic consortia, and, in the absence of sulphate, through CO_2_ reduction to methane by ANME-2. Under these conditions, the production of methane was one ninth the rate of methanotrophy coupled to sulphate-reduction. Paired single cell FISH-nanoSIMS analysis of anabolic activity indicates that CO respiration appears to support cell maintenance rather than active growth, consistent with the observed down-regulation of energy generating complexes in ANME (e.g., *mtr*, *rnf*, etc.). The versatile capability of CO oxidation by anaerobic methanotrophic consortia broadens our understanding of carbon cycling in methane seeps and highlights potential mechanisms of resilience by methanotrophic archaea under changing geochemical regimes.

## Introduction

The anaerobic oxidation of methane (AOM) coupled to sulphate reduction is an important microbial process for controlling methane flux from ocean sediments worldwide (*1*, *2*). This process is largely mediated by anaerobic methanotrophic archaea (ANME) living syntrophically in consortia with sulphate-reducing bacteria (SRB) (*3–5*). Methanotrophic ANME-2 archaea (*Ca.* Methanogastraceae, Methanocomedens, and Methanomarinus) (*6*) in ocean sediments have been shown to be energetically versatile, capable of using terminal electron acceptors besides sulphate, including metal oxides (manganese (birnessite) and iron (ferrihydrite)) (*7*), and humic acid-like substances (*8*). However, the potential for these archaeal methanotrophs to oxidize terminal electron donors other than methane, has not been specifically investigated, beyond experiments assessing possible syntrophic intermediates for SRB in the absence of methane (*9*).

Members of the ANME-2 within the order *Methanosarcinales* are evolutionarily related to methylotrophic methanogens such as *Methanosarcina* (*6*). Previous studies showed that *Methanosarcina* species were able to grow on carbon monoxide (CO) as an energy source (*10*, *11*), suggesting the possibility that related members of ANME-2 may also utilize CO. While infrequently measured in geochemical investigations of seep environments, some reports indicate that CO can be produced in sediments, resulting from the thermal decomposition of organic compounds such as humic acids and phenolic compounds or released as a by-product from microbial pathways (*12–19*). From a thermodynamic perspective, CO represents a favourable electron donor for ANME with a low reduction potential of –558 mV (pH 7.0, CO_2_/CO) (*20*) and there are several examples of CO oxidizing microorganisms within diverse environments. This includes anaerobic bacteria such as *Moorella thermoacetica* (*21*, *22*), the purple sulphur bacteria *Rhodopseudomonas rubrum* (*23*, *24*), *Carboxydothermus hydrogenoformans* (*25*), selected sulphate-reducing bacteria (*26*), as well as archaea (*Candidatus Hydrothermarchaeota)* living within the subseafloor crust (*27*). Metagenomic surveys from geographically diverse methane cold seeps have additionally showed widespread distribution of anaerobic CO dehydrogenases (CODH) among diverse ANME lineages including the mono-functional CODH and bi-functional CODH involved in the Wood–Ljungdahl pathway, supporting the potential for CO metabolism by ANME.

This study investigated whether environmental methanotrophic archaea belonging to the ANME-2, and their syntrophic SRB partners, can metabolize CO using a combination of anoxic microcosm experiments, geochemical and isotopic analysis, metatranscriptomics, and single-cell resolved fluorescence *in situ* hybridization and nanoscale secondary ion mass spectrometry (FISH– nanoSIMS).

## Results

### Both ANME and SRB encode carbon monoxide dehydrogenases

Anaerobic CO dehydrogenases (CODHs) catalyzing the conversion of CO and CO_2_ are comprised of two main groups, the CO dehydrogenase/acetyl-CoA synthase complex (*cdhA*), present in the Wood–Ljungdahl pathway, and the monofunctional anaerobic CO dehydrogenase (*cooS*) (*28*). To examine the distribution and diversity of CO dehydrogenases in ANME-SRB consortia, we conducted a phylogenetic analysis of CO dehydrogenases within representative high-quality genomes for all described marine ANME groups including *Candidatus* Methanophaga (ANME-1), *Ca.* Methanocomedens (ANME-2a), *Ca.* Methanomarinus (ANME-2b), *Ca.* Methanogaster (ANME-2c), and *Ca.* Methanovorans (ANME-3) (*6*); and syntrophic SRB lineages (*Ca.* Syntrophophila gen. nov., (Seep-SRB1a), *Ca.* Desulfomellonium gen. nov., (Seep-SRB1g), *Ca.* Desulfomithrium gen. nov. (Seep-SRB2), and *Ca.* Desulfofervidus sp. (HotSeep-1 cluster)) (*29*), representing a total of 49 ANME and 65 SRB MAGs. Both *cdhA* and *cooS* were detected in ANME and SRB lineages, with the exception of ANME-1 representatives that were lacking *cooS* homologs (Fig. 1, Extended Data Fig. 1). In addition, several MAGs, including ANME-2a CONS7142G09b1 and Seep-SRB1g str. CR10073A, were found to host two copies of *cooS.* Both of these were verified to occur on contigs with other characterized ANME-2a or Seep-SRB1g genes and unlikely to be a product of mis-binning (Fig. 1). These observations illustrate the broad distribution of CO dehydrogenases within diverse ANME and SRB lineages. This genomic evidence motivated our deeper physiological study of the potential for CO metabolism by anaerobic methanotrophic consortia.

**Fig. 1.**
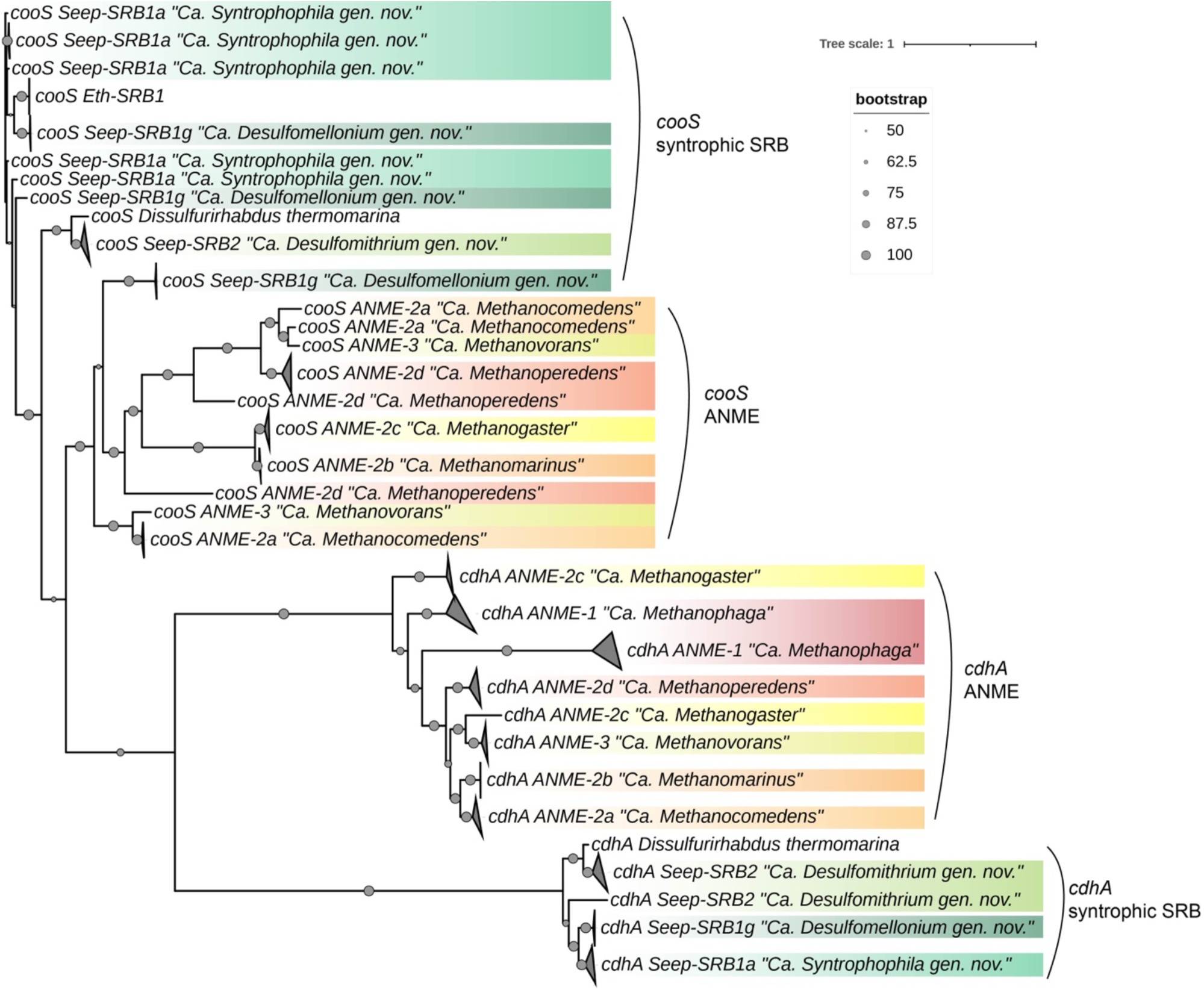
The phylogeny of CO dehydrogenase/acetyl-CoA synthase (CODH/ACS) complex alpha subunit (*cdhA*) and monofunctional anaerobic CO dehydrogenase (*cooS*) in ANME and syntrophic SRB lineages. The CODH homologs were retrieved from metagenome assembled genomes reported in previous studies (*6*, *29*). All homolog alignments were examined manually after identification with BLAST. We operationally distinguished *cdhA* from *cooS* homologs by the following principle: a hit was labelled *cdhA* if it was part of a CODH/ACS complex cluster (operon), and *cooS* if not in the CODH/ACS operon.

### ANME-SRB consortia can respire CO and sulphate

To test whether ANME-SRB consortia can respire CO, we performed a series of anaerobic microcosm experiments using deep-sea methane seep sediments collected from the Costa Rica margin (see *Methods*). The sediments were dominated by members of *Methanomarinus* (ANME-2b; relative abundance 53.63%) along with *Methanocomedans* ANME-2a-2b (9.12%) and *Methanogaster* (ANME-2c; 2.75%) groups, and their sulphate-reducing bacterial partners Seep-SRB1 (6.21%) based on 16S rRNA amplicon sequencing (Extended Data Fig. 2).

We tested the ability to oxidize CO coupled with sulphate using ^13^CO at three different concentrations, 0.1-, 0.4-, and 1.0-bar partial pressure in biological triplicates. A positive control (methane + sulphate; n = 3) and a killed control (n = 1; 0.1-bar ^13^CO + sulphate) were also run in parallel (Supplementary Table 1). Labelled ^13^CO was used to quantify the rate of CO oxidation by quantifying the increase of ^13^C-DIC over time concomitant with the reduction of sulphate to sulphide (HS^−^) over the course of 24 days. These experiments confirmed active CO oxidation alongside the reduction of sulphate, however CO oxidation rates showed a decrease with increasing CO partial pressures (Fig. 2). At 0.1 bar, CO was oxidized at a rate of 34.97 ± 0.68 µM d^−1^ cm^−3^_sed_, equating to approximately half the rate of methane oxidation in parallel control incubations with ^13^CH_4_ (64.63 ± 0.68 µM d^−1^ cm^−3^_sed_); (Supplementary Table 2). This rate decreased further from 23.21 ± 4.67 µM d^−1^ cm^−3^_sed_ to 12.00 ± 0.07 µM d^−1^ cm^−3^_sed_ with headspace CO concentrations of 0.4 bar and 1.0 bar, respectively (Supplementary Table 2), possibly attributed to CO toxicity at high concentrations as reported in other organisms (*10*, *30*).

**Fig. 2.**
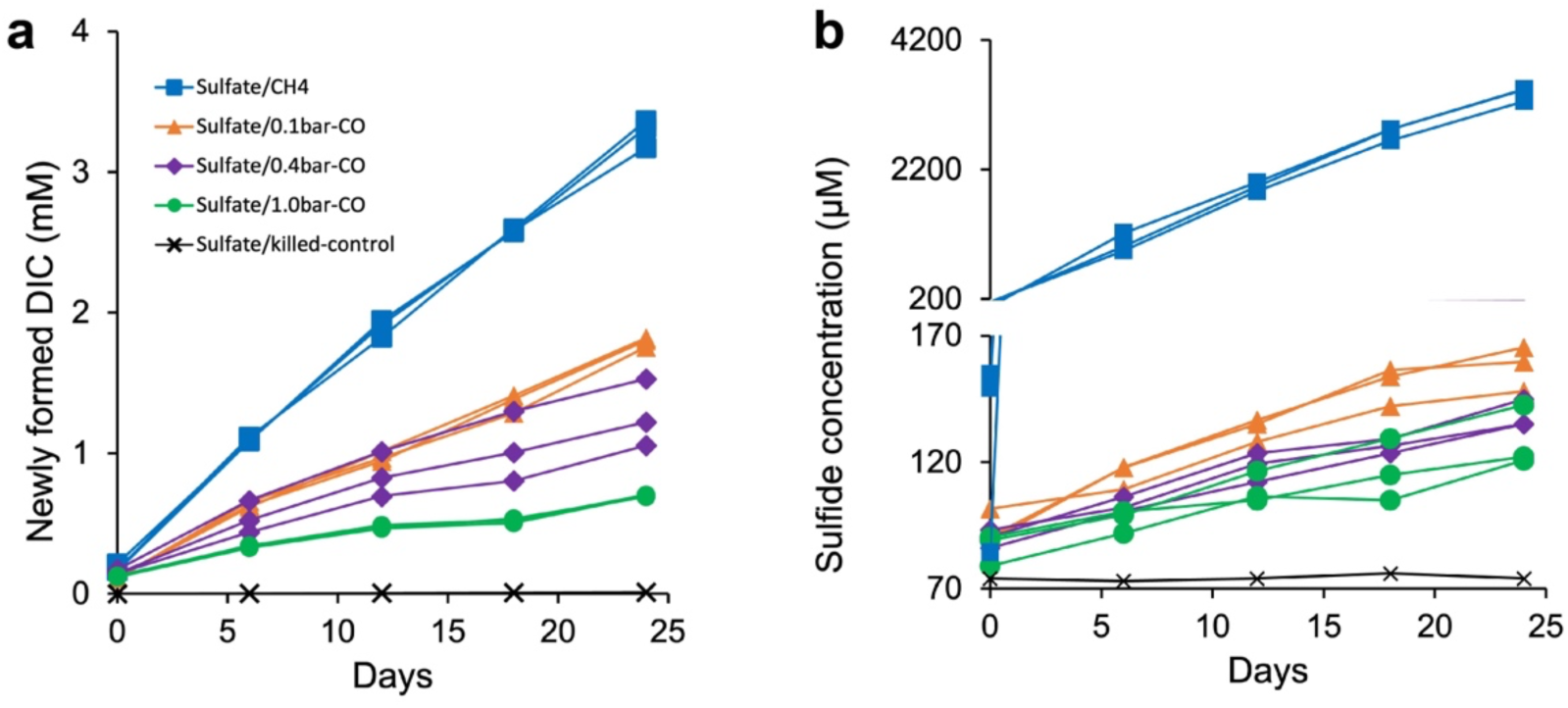
Time course of CO and methane oxidation by anaerobic methanotrophic consortia within methane seep microcosm experiments containing either CH_4_ or CO as the electron donor and sulphate as the electron acceptor. **a**, The oxidation of ^13^CH_4_ or ^13^CO to ^13^C-dissolved inorganic carbon (DIC); **b**, the reduction of sulphate reflected by the production of hydrogen sulphide (HS^−^).

### CO oxidation includes an additional electron sink besides sulphate

An examination of the electron transfer and mass balance for the CO and sulphate redox reaction indicates that there were significantly higher rates of CO oxidation in the incubation experiments than could be explained by sulphate reduction alone. Indeed, the measured rate of CO_2_ production (34.97 µM d^−1^ cm^−3^_sed_) was 7 times greater than the predicted rate (5.32 µM d^−1^ cm^−3^_sed_) calculated from sulphide production based on Eq. 1 (Supplementary Table 2). This discrepancy suggests that, under conditions with CO and sulphate, sulphate accounts for only a portion of the electrons from CO oxidation, implying that there must be additional electron acceptor(s) in the sediment microcosm beyond sulphate.

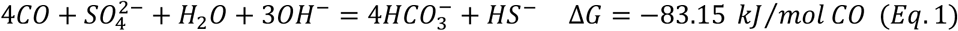

Based on this, we identified two likely alternative electron acceptors, CO and CO_2_ (present as bicarbonate (HCO_3_^−^) in the medium), that were in high enough concentrations to account for this discrepancy. First, CO itself could be reduced by nitrogenases present in ANME and/or SRB (*31*, *32*). Electron carrier ferredoxin, being reduced at CO dehydrogenases, could feed the reduction of CO by nitrogenases with an investment of ATP and subsequently result in the production of hydrocarbons (see illustration in Extended Data Fig. 3). Previous work demonstrated that both vanadium- and molybdenum-nitrogenases are able to reduce CO to diverse hydrocarbons including ethylene (C_2_H_4_), ethane (C_2_H_6_) and propane (C_3_H_8_) (*33*, *34*). However, we did not detect any of these alkanes when we analyzed our microcosm experiments using GC-MS (data not shown) and we therefore presume that any potential CO reduction by ANME or SRB nitrogenase is minimal to non-existent.

Another possibility is coupling CO with CO_2_ (HCO_3_^−^) as the electron acceptor, a reaction known to be catalysed by *M. acetivorans*. In this study, a number of products were reported including formate, acetate and methane (*10*). Neither formate nor acetate were detected by either NMR or IC in our microcosm experiments (detection limits: formate on NMR, *ca.* 3 µM; acetate on IC, *ca.* 5 µM; data not shown). We then focused on methane production. This reaction is predicted to be thermodynamically favourable, with the reduction potential of CO_2_/CH_4_ (−245 mV at pH 7.0) (*8*) higher than that of CO_2_/CO (−558 mV, pH 7.0). Microcosm incubations amended with 0.1 bar CO and 5 mM sulphate confirmed that CO continued to be oxidized over time alongside sulphate reduction (Fig. 3a, b). These incubations showed evidence of methane production however the concentration did not accumulate above *ca.* 0.2 µmol in the headspace (Fig. 3c). We hypothesized that this trace methane concentration, lower than predicted, resulted from subsequent oxidation via sulphate-coupled AOM (i.e., cryptic methane cycling) (*35*). To test this idea, we conducted follow up microcosm experiments amended with ^13^CH_4_ and SO_4_^2-^, supporting conventional sulphate-coupled AOM, and then spiked in unlabelled CO (^12^CO) at day 40 to track the potential contribution of CO-derived CH_4_ above that of ^13^CH_4_ to AOM. Newly formed ^13^C- and ^12^C-DIC quantification was used to differentiate the relative proportion of CH_4_ oxidation and CO oxidation activity, assuming ^12^C-DIC generation after day 40 was attributed to CO. These results supported the active oxidation of methane by ANME-SRB consortia in the presence of CO (Fig. 4).

**Fig. 3.**
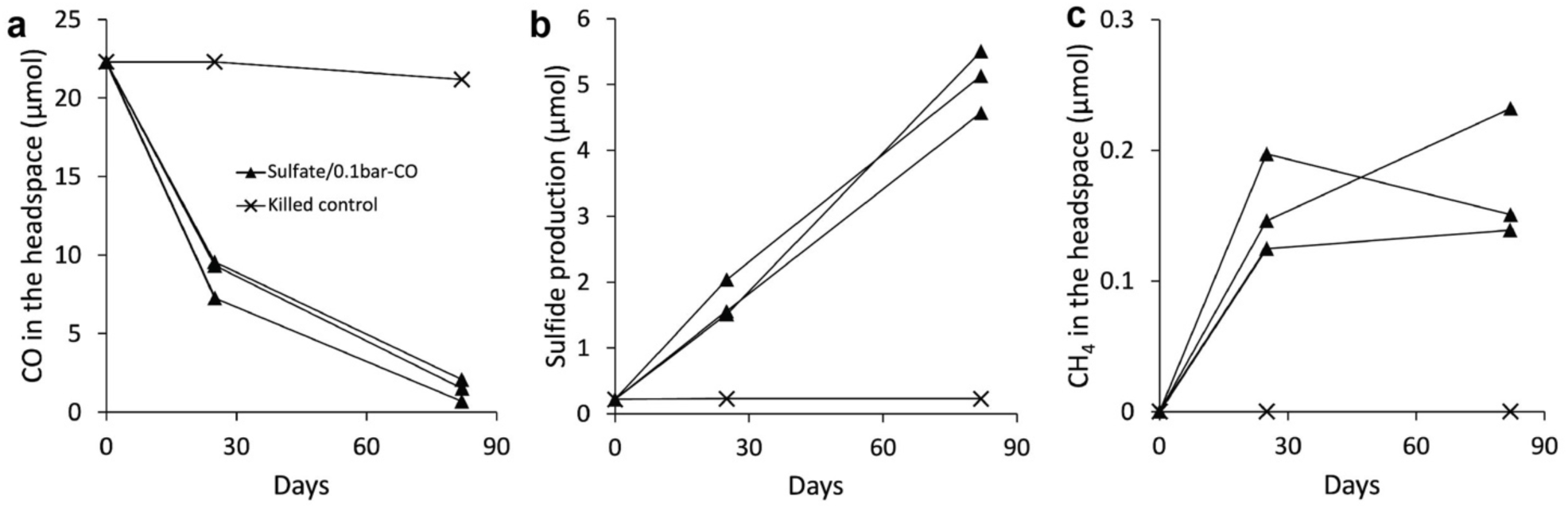
CO oxidation and corresponding sulphide and methane production monitored over time from microcosm incubations amended with sulphate. Biological triplicates were used for each experimental condition and shown separately along with a single killed control incubation (× symbol).

**Fig. 4.**
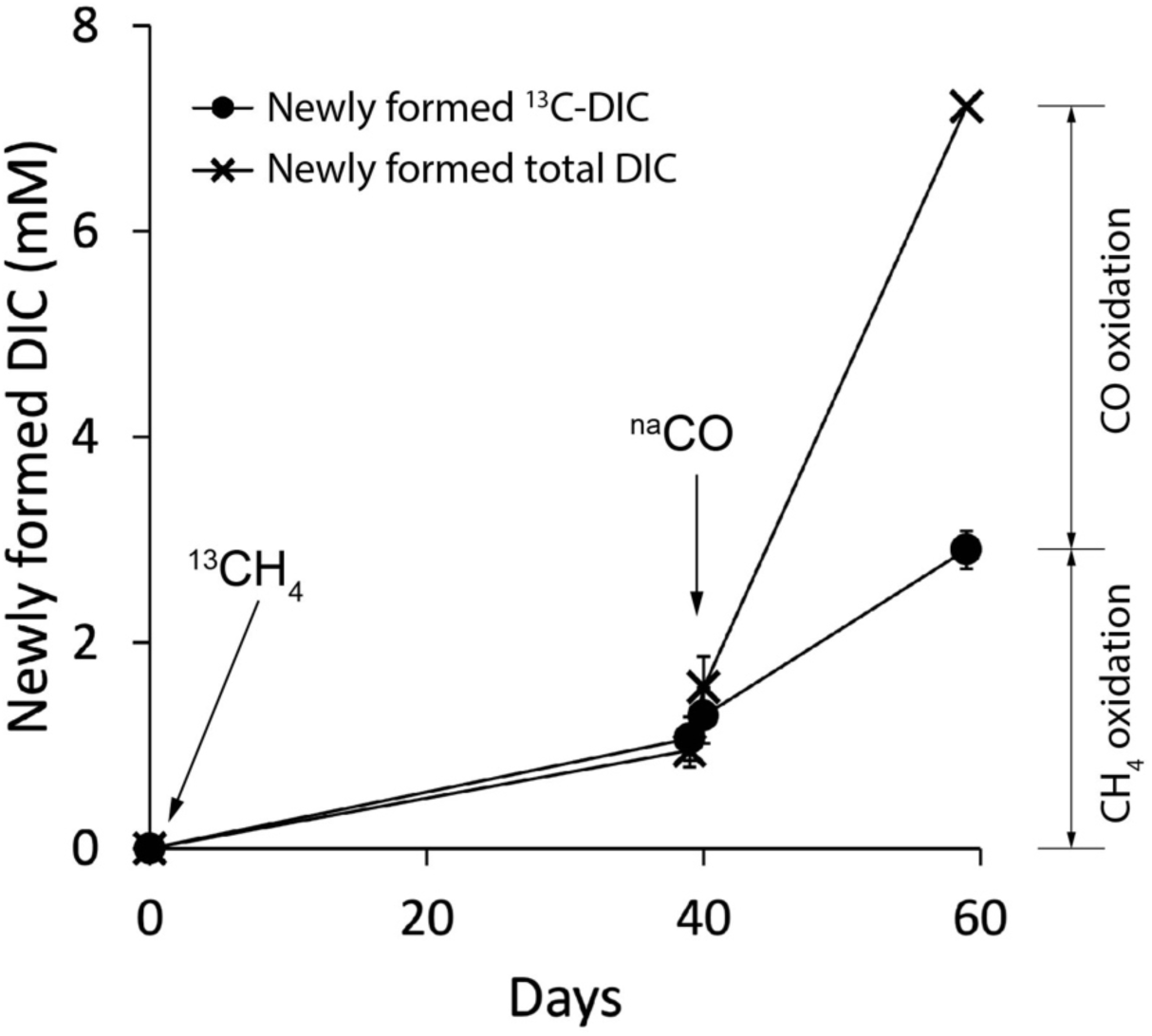
DIC production in microcosm experiments containing sulphate, methane and CO. Microcosms were initially amended with ^13^CH_4_ and sulphate. ^12^C- and ^13^C-DIC comprised of the total newly formed DIC, where ^13^C-DIC came from ^13^CH_4_ oxidation. Unlabelled CO (^na^CO, 0.4 bar) was then added at day 40 of the experiment to examine the proportion of DIC sourced from methane vs. CO. A mix of ^13^CH_4_ and ^na^CO were both present in the headspace after day 40 and served as potential electron donors. Subtracting the ^13^C-DIC from the total concentration of newly formed DIC after day 60, differentiated the proportion of CO oxidation and methane oxidation occurring. This experiment was performed in biological triplicate (mean ± s.d., n = 3). Where not visible, the error bar is smaller than the symbol.

To further investigate the potential for methane production from CO oxidation in the absence of sulphate-coupled AOM, we conducted a second set of microcosm experiments under sulphate-free conditions and amended with CO (0.4 bar partial pressure; Supplementary Table 1). These incubation conditions limit the activity of the syntrophic SRB, minimizing any subsequent oxidation of produced CH_4_ through sulphate-coupled AOM. By supplying HCO_3_^−^ as the sole added terminal electron acceptor, a continuously increasing amount of methane was produced coupled to CO consumption at a rate of 0.36 µmol CO d^−1^ cm^−3^_sed_ (mean, n = 2; Fig. 5). Parallel microcosm experiments without the addition of HCO_3_^−^ to the medium were also run. These results showed CO oxidation was also coupled with methane production under these conditions and at comparable rates of 0.35 µmol CO d^−1^ cm^−3^_sed_ (mean, n = 3). This suggests that self-generated CO_2_ may also serve the terminal electron acceptor. Most importantly, the ratio of CO oxidation rate to methane production rate calculated from the five replicates was 3.85 ± 1.93, close to the predicted stoichiometry of 4:1 in Eq. 2.

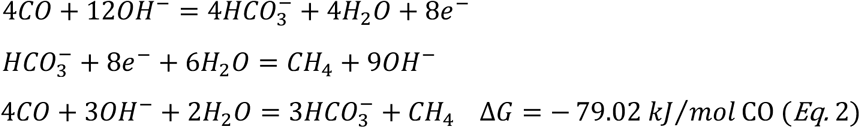

**Fig. 5.**
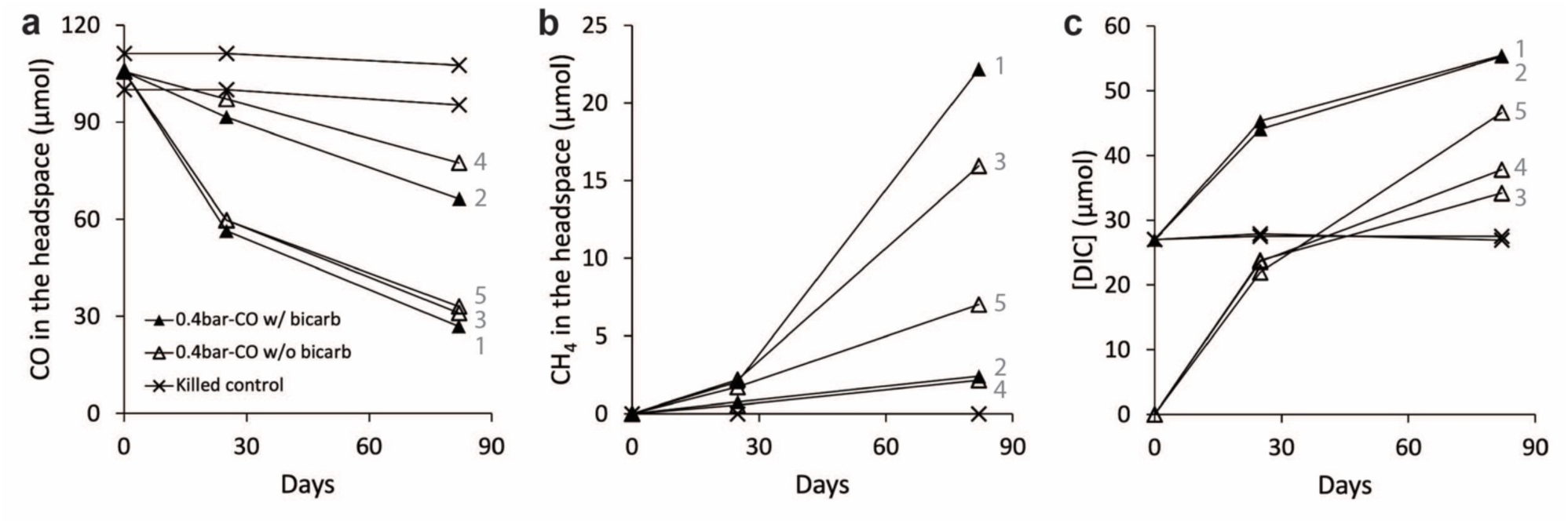
CO consumption coupled to methane production over time in microcosms without sulphate and with bicarbonate as the sole added terminal electron acceptor. Biological duplicates were performed in microcosms with bicarbonate as the sole electron acceptor (solid triangles). Biological triplicates were run without the addition of an external electron acceptor (open triangles). A killed control in duplicate was also run (× symbol). **a,** Changes in CO concentrations over time; **b,** corresponding methane generation over time. Units were normalized to the total molar amount as µmol. The numbers to the right of each line correspond to individual bottles in this experiment.

To confirm that methane was sourced from CO_2_ in these sulphate-free incubations, experiments amended with ^13^CO (≥ 99 atom % ^13^C) and ^12^C-HCO_3_^−^ (100%) were used (incubation set #4; Supplementary Table 1). In these experiments, the detection of ^13^C in the bicarbonate pool is predicted to occur through CO oxidation (Eq. 2; Fig. 5c), and any subsequent methane production from CO_2_ reduction will show a mixed ^12^C/^13^C ratio if both processes are occurring (Eq. 3). Results from these isotope labelling experiments were consistent with this hypothesis, producing partially ^13^C-enriched CH_4_ with a ^13^C value of 40.64 ± 0.68% (n = 3); (Extended Data Fig. 4).

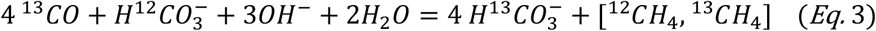

### ANME and SRB consortia show anabolic activity by FISH–nanoSIMS when supplied with CO as the electron donor

Having demonstrated that CO oxidation occurs in seep sediments coupled with the reduction of either sulphate or CO_2_, we then asked whether this bulk anaerobic CO respiration in turn supported anabolic activity by ANME-2 and their SRB partners in the absence of methane addition. Single cell anabolic activity was measured using stable isotope probing with ^15^N-ammonium as a marker for biosynthetic activity combined with FISH–nanoSIMS (*36–38*). In these 4-month CO incubations, sulphate and bicarbonate served as electron acceptors and ^15^N-ammonium was added as a nitrogen source (Supplementary Table 1).

A total of 4,979 FISH-identified cells within 42 multi-celled consortia of ANME-2b (ANME-2b-729 oligonucleotide probe) and their SRB partner (likely Seep-SRB1g (*32*); hybridized with a *Desulfosarcina/Desulfococcus* group probe (DSS658)) were analysed from the Set #1 incubation experiments (Supplementary Table 1). Cellular ^15^N enrichment was measured in both ANME-2b and syntrophic SRB cells (Fig. 6a and 6b, Table 1), demonstrating that both were anabolically active during CO oxidation coupled with sulphate reduction in the absence of external methane addition. Notably, high concentrations of CO in the headspace resulted in lower levels of cellular ^15^N enrichment, consistent with some degree of toxicity (Fig. 6b, Table 1). Anabolic activity decreased to a greater extent with increasing concentrations of CO for both ANME (1.0 bar CO) and SRB (0.4 bar and 1.0 bar), with values below the baseline ^15^N enrichment (here determined to be 0.53 atom %; Table 1). This observation is consistent with reports from CO metabolizing methanogens, where higher levels of CO inhibited methanogenic activity (*10*, *39*). When CO concentrations were provided at 0.4 bar, however, approximately 22% of the 831 ANME-2b cells analysed remained anabolically active at a considerable level, while the fraction of active SRB cells dropped dramatically (6 out of 488 cells). This suggests a higher CO tolerance by ANME-2b relative to their SRB partner.

**Fig. 6.**
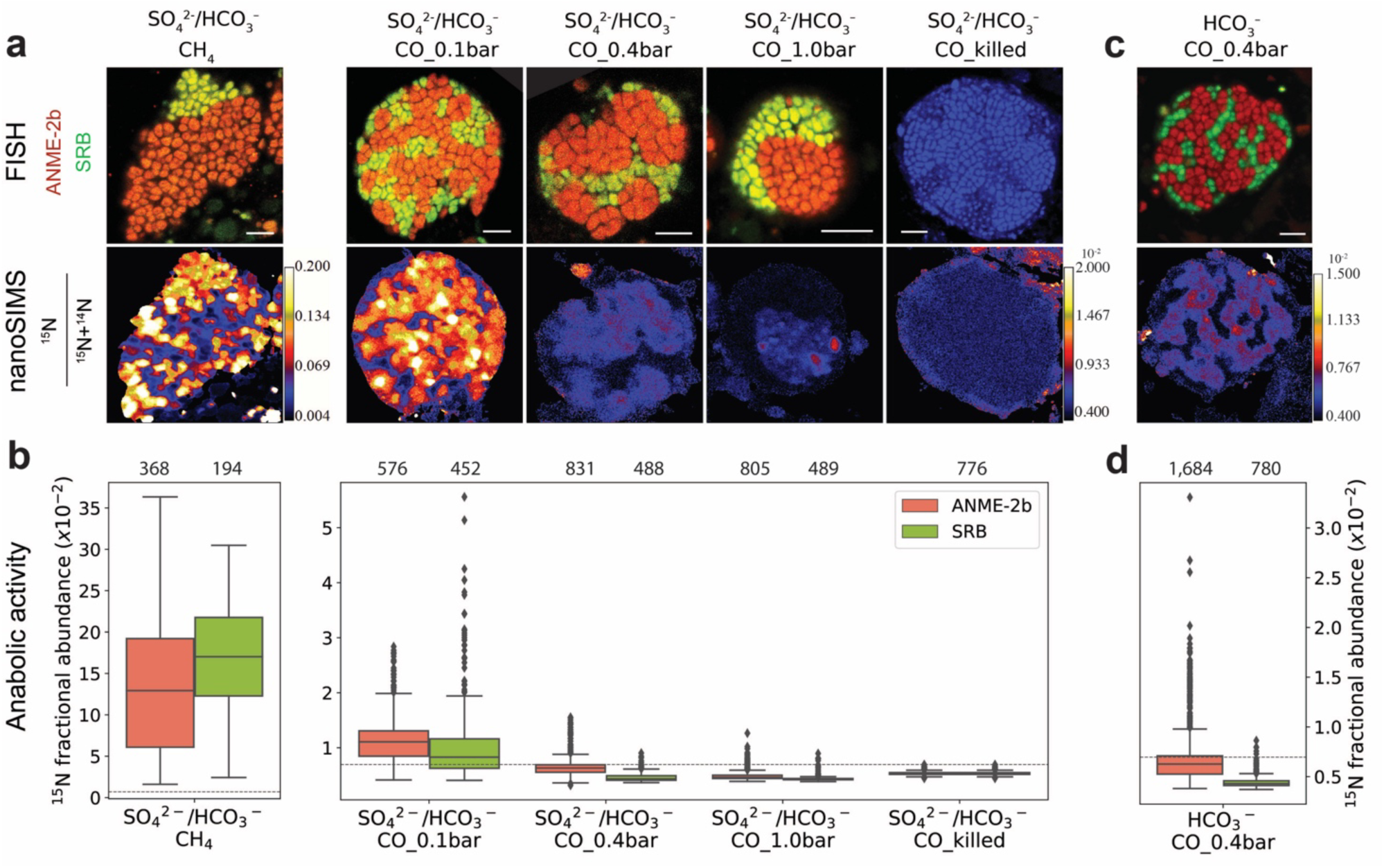
Anabolic activity summarized from paired FISH–nanoSIMS experiments for ANME-2b archaea and its syntrophic SRB (SeepSRB1g) In panel **a** and **c**, the top row shows FISH images for ANME-2b (red) and their syntrophic SRB partner (green); the bottom shows a paired representative nanoSIMS image for each microcosm condition. The ^15^N fractional abundance in all cell ROIs analysed in this study is summarized in panels **b** and **d**, where the dashed line represents the minimum threshold considered for cell ^15^N enrichment/activity (0.59 atom% ^15^N). Statistical information for these analyses is summarized in Table 1. (**a**, **b**) Samples from sulphate/HCO_3_-microcosm incubations were ended after 122-days, and (**c**, **d**) sample from HCO_3_^−^ microcosms (sulphate free) were ended after 90-days. The numbers above (**b**) and (**d**) represent the number of ANME and SRB cells (ROIs) analysed in each treatment. Scale bar is 5 µm in all panels.

**Table 1.**
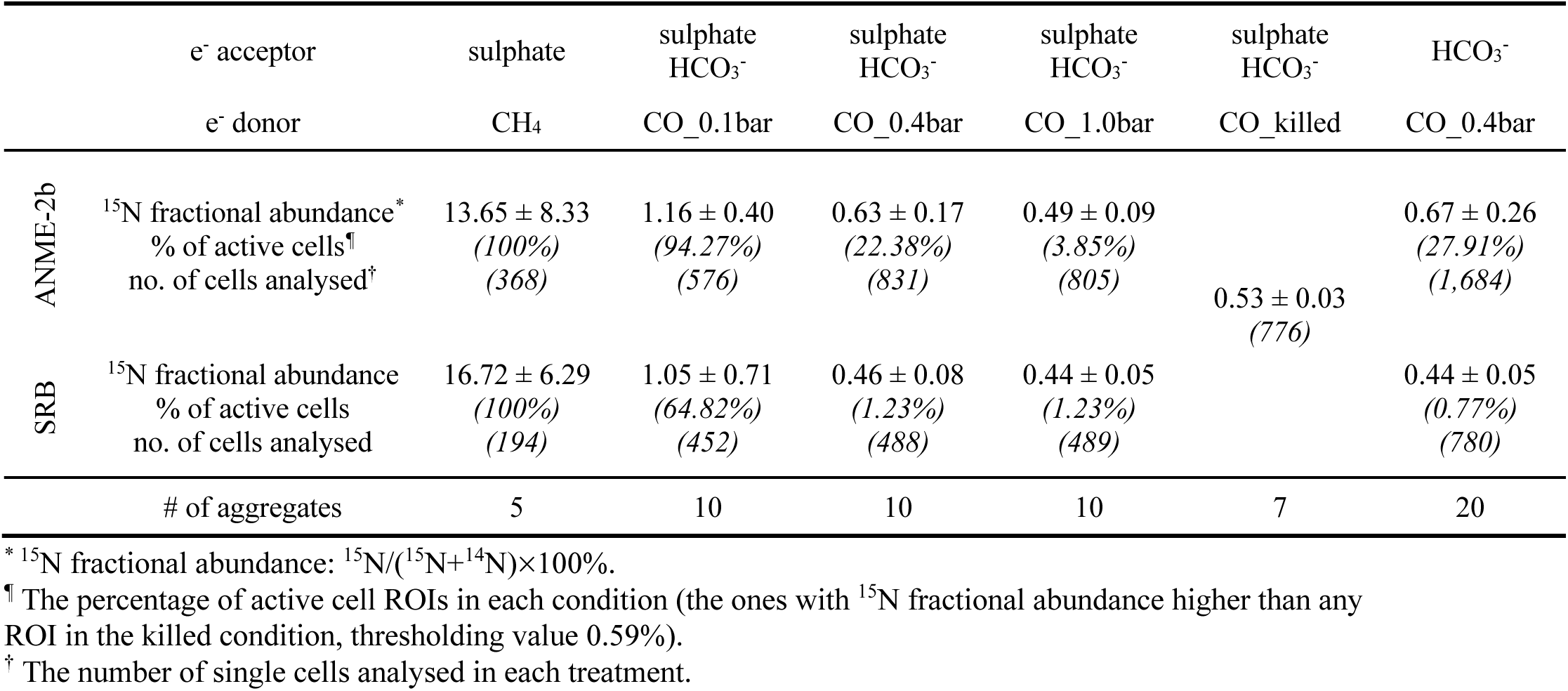
^15^N enrichment of ANME-SRB consortia at single cell level from FISH–nanoSIMS analysis.

To further examine whether ANME-2b were anabolically active and responsible for the measured CO oxidation and methane production rather than catalysed by other unidentified CO-oxidizing microorganisms in the sediment, additional FISH–nanoSIMS experiments were performed using sulphate-free conditions over a 3-month period, here designed to limit the activity of the SRB partner and potential for cryptic methane cycling through syntrophic AOM (Fig. 6c and 6d, Table 1). Cellular ^15^N enrichment for 2,464 ROIs (cells) belonging to ANME-2b and their SRB partners were resolved. ANME-2b cells showed an average ^15^N enrichment of 0.67 ± 0.26 atom % (470 out of 1,684), while co-occurring SRB cells were anabolically inactive as expected, with the exception of a few cells (6 out of 780); (Fig. 6d). The observation of ANME-2b anabolic activity in the absence of activity by their SRB partner supported their direct involvement in methane production during respiration of CO as an energy source.

We assessed the relative differences between CO oxidation and anaerobic methanotrophy supporting ANME growth using the cell-specific growth rates from FISH-nanoSIMS and bulk sediment respiration rates (Table 2). With CO addition (0.4 bar), methane production by ANME-2b occurred with an average rate that was approximately 1/9 of the sulphate-coupled AOM rate determined in parallel incubations supplied with methane and sulphate (Table 2). However, the apparent cellular energy gain from growth rate was minimal, with ANME-2b growth on CO (0.4 bar) and CO_2_ nearly 20 times lower than the canonical AOM coupled sulphate reduction (Table 2). This implies that the observed ANME-2b CO oxidation is likely used to support cellular maintenance processes, rather than growth.

**Table 2.**
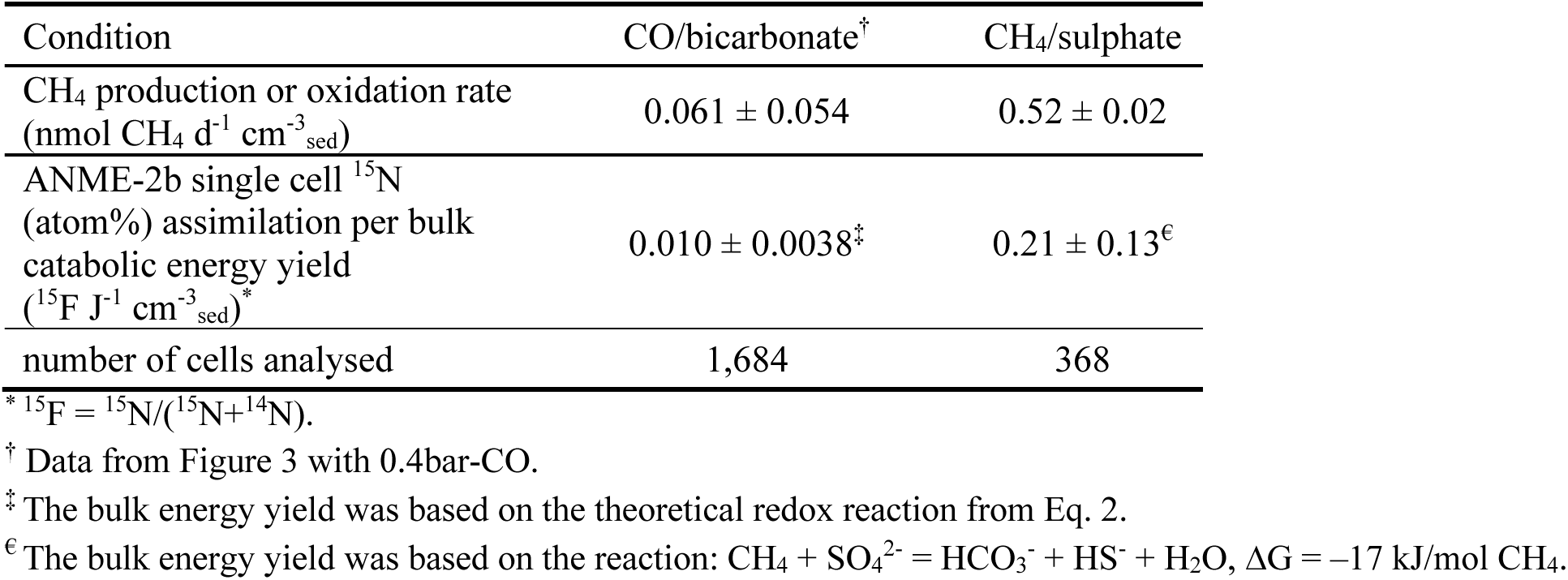
The ratio of single cell ^15^N assimilation to bulk potential energy generation by redox reaction.

### Metatranscriptomic analysis of ANME-2b/SRB response to CO

Metatranscriptomics was performed to investigate the underlying metabolic response of CO metabolism compared with methane (1.5 bar methane, 0.1 and 1.0 bar CO) using either sulphate or bicarbonate as the electron acceptors (n = 3 biological triplicates for each treatment, 9 sediment incubations total). Nearly 90% of the reads from these microcosm experiments were affiliated with the dominant AOM consortia members, ANME-2b and syntrophic SeepSRB-1g, consistent with the 16S rRNA amplicon sequencing results (Extended Data Fig. 2). In the remaining pool of transcripts, we were not able to resolve the taxonomic identity of any other microbes at a diagnostic level, however targeted screening of expressed methyl coenzyme M reductase (*mcr*) from methanogens only recovered ANME-affiliated *mcr* transcripts. This finding further decreased the likelihood of hidden CO-metabolizing methanogens in the sediment incubations. The dominance of transcripts affiliated with ANME-SRB consortia members supports our single cell isotopic data and independently supports ANME-2b and their syntrophic SRB partner as the organisms responsible for the observed CO oxidation. Subsequent transcriptomic analyses were then focused on ANME-2b and Seep-SRB1g. To better illustrate the transcriptomic activity, both differential gene expression (hereafter abbreviated DE) (DESeq2) and percentile ranking (Sleuth TPMs (transcripts per million reads)) were analysed.

For ANME-2b, the expression of most genes did not change significantly between CO and methane treatments according to the overall transcripts per million reads (TPM) distribution (Extended Data Fig. 6). Approximately 2.79% of ANME genes were significantly up-regulated and 13.21% down-regulated in the microcosms with 0.1 bar CO compared with the control AOM conditions supplied with CH_4_ and sulphate (Extended Data Fig. 7). Of the genes found to be significantly up-regulated in the CO condition compared to the CH_4_ condition, most were associated with genes generally affiliated with stress response, including defence mechanisms, toxin/antitoxin systems, and stress-related ABC exporters (ABC-type antimicrobial peptide transport system); (Supplementary Data 1). This indicates that CO is a less favourable substrate for ANME-2b respiration compared with CH_4_ and consistent with some degree of toxicity.

The ANME-2b carbon monoxide dehydrogenase, *cooS,* was actively expressed consistent with involvement in CO metabolism. Unexpectedly however, the expression levels were found to be equivalent between the methane/sulphate vs. CO/sulphate or CO/bicarbonate conditions, with mean values of 52.3 ± 5.1% (CH_4_), 48.2 ± 3.4% (0.1 bar CO), and 55.1 ± 3.4% (1.0 bar CO), respectively (Supplementary Data 1). In contrast to *cooS*, significantly lower expression levels were found for the full *cdh* operon, which is an integral component of the Wood-Ljungdahl pathway (66.3 ± 13.6% (CH_4_) vs. 49.9 ± 11.6% (0.1 bar CO), *p* < 0.001 (two-tailed *t*-test)); (Supplementary Data 1). This suggests a different response between the two types of ANME-2b-encoded CO dehydrogenases when supplied with an external source of CO.

For ANME-2b, we observed a high level of expression for most genes associated with methanotrophy/methanogenesis and carbon fixation pathways within the CO/sulphate and bicarbonate condition, that was at a comparable level with the methane/sulphate treatment supporting AOM (Fig. 7, Supplementary Data 1). These genes included the methyl-coenzyme M reductase (*mcr*) operon (>99.8% among all conditions), 5,10-methylenetetrahydromethanopterin reductase (*mer*); (mean: 99.1% (CH_4_), 97.4% (0.1 bar CO), 96.8% (1.0 bar CO)), and formylmethanofuran dehydrogenase (*fwd*) operon (mean: 96.6% (CH_4_), 94.0% (0.1 bar CO), 93.8% (1.0 bar CO)); (Fig. 7, Supplementary Data 1). In contrast, N^5^-methyl-H_4_SPT:HS-CoM methyltransferase (*mtr*) operon, a key energy-converting step in the methanogenesis pathway, had a significantly lower expression level in CO/sulphate/bicarbonate conditions compared to the parallel methane/sulphate treatment (0.1 bar CO vs. CH_4_; percentile: 84.7 ± 3.3% vs. 98.2 ± 0.6%, mean ± s.d.; differential analysis: *ca.* 25-fold decrease on average); (Fig. 7, Extended Data Fig. 8, Supplementary Data 1).

**Fig. 7.**
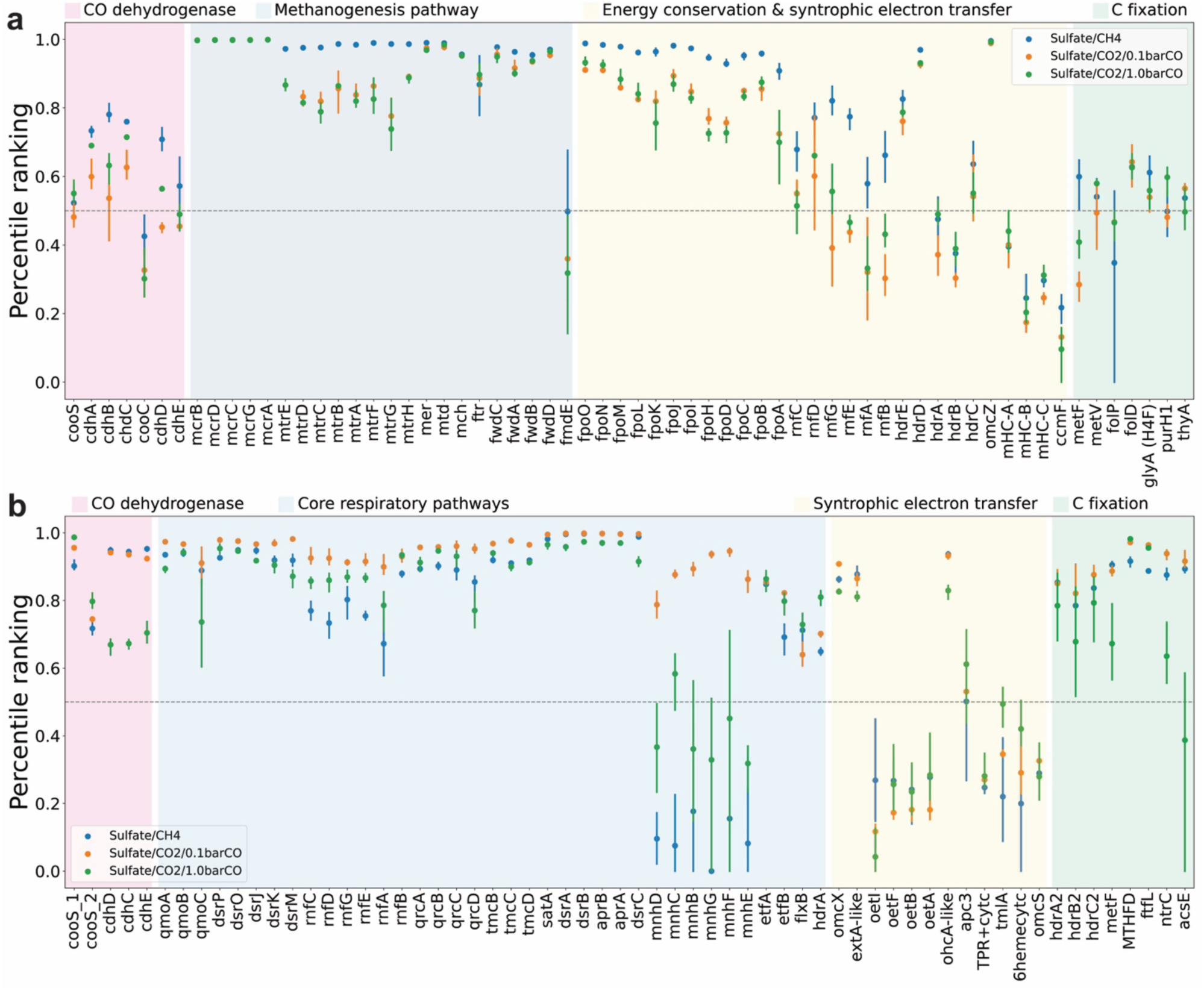
Transcriptomic data for selected genes from ANME-2b and syntrophic SeepSRB-1g involved in energy metabolism. Transcriptomic analysis for seep sediment microcosms were conducted after 6 days of incubation with either methane or CO as the electron donor and sulphate as the electron acceptor. **a**, Expression of selected genes associated with ANME-2b: CO dehydrogenase, methanogenesis pathway, energy conservation and syntrophic electron transfer, and carbon fixation. **b,** Expression of selected genes in SeepSRB-1g: CO dehydrogenases, core respiratory pathways, syntrophic electron transfer, and carbon fixation. The detailed profiling of these genes are summarized in Supplementary Data 1 and 2. Symbol colours here correspond to the microcosms shown in Fig. 2.

Other key genes involved in the pathways of energy conservation and syntrophic electron transfer (*6*) also displayed a lower expression level in treatments with CO versus methane, both in terms of ranking and DE analysis (Fig. 7, Extended Data Fig. 8). For example, the NADH-ubiquinone/plastoquinone oxidoreductase (*fpo*) operon (F_420_H_2_:methanophenazine oxidoreductase), involved in F_420_ conversion, was down-regulated under CO/sulphate/bicarbonate treatments (ranking: 84.9% (0.1 bar CO) and 83.9% (1.0 bar CO) vs. 96.7% (CH_4_)), with a 9-fold decrease in differential expression (DE) (0.1 bar CO vs. CH_4_; Fig. 7, Extended Data Fig. 8, Supplementary Data 1). In addition, the expression of genes associated with the Na^+^-translocating ferredoxin:NAD^+^ oxidoreductase (*rnf*) was also significantly lower in CO/sulphate/bicarbonate treatments (ranking: mean, 71.4% (CH_4_), 43.4% (0.1 bar CO), 49.4% (1.0 bar CO); DE: *ca.* 4-fold decrease (0.1 bar CO vs. CH_4_); Fig. 7, Extended Data Fig. 8, Supplementary Data 1). The decrease in transcription of *mtr*, *fpo*, and *rnf* indicates that while these energy conserving pathways are expressed during CO oxidation in ANME-2b, this respiratory process clearly deviates from their primary methanotrophic metabolism.

In comparison with ANME-2b, the syntrophic SeepSRB-1g partner revealed an opposite transcriptional response to CO, where 21.00% of expressed genes were significantly up-regulated in treatments with 0.1bar CO with either sulphate or bicarbonate as the electron acceptor compared with the methane/sulphate treatment (Extended Data Fig. 6), and only 1.52% of genes were significantly down-regulated (Extended Data Fig. 7). This expression pattern changed substantially in the high CO condition (1.0 bar CO), where the proportion of down-regulated genes increased to 14.42%, and only 2.32% of genes were highly expressed (Extended Data Fig. 7). Like ANME-2b, SeepSRB-1g actively expressed CO dehydrogenases among CH_4_ and CO treatments, where both the *cooS* and the *cdh* operons had comparable rankings in expression between methane/sulphate and CO/sulphate/bicarbonate (0.1 bar CO) condition (Fig. 7, Supplementary Data 2). At higher CO concentrations however, *cdh* operon gene expression was significantly down-regulated (1.0 bar CO).

In the CO incubations with sulphate, the Seep-SRB1g partner showed active expression of respiratory pathways, potentially enabling DIET between the syntrophic partners (*38*). This included significant up-regulation of genes encoding extracellular multi-haem cytochrome *c* proteins predicted to be involved in extracellular electron transfer (*29*) including *omcX* (fold change: 2.72 (1.0bar-CO *vs.* CH_4_), *P_adj_* = 5.78E-08) and *tmlA* (fold change: 3.15 (1.0bar-CO vs. CH_4_), *P_adj_* = 0.027); (Extended Data Fig. 8, Supplementary Data 2). Most genes involved in the dissimilatory sulphate-reduction pathway were also highly expressed in all three conditions (percentile ranking >80%) including the quinone-interacting membrane-bound oxidoreductase complex (*qmo*), dissimilatory-type sulphite reductase (*dsr*), sulphate adenylyltransferase (*sat*) and adenylyl-sulphate reductase (*apr*); (Fig. 7, Supplementary Data 2). In addition, the expression level of genes belonging to the multi-subunit Na^+^/H^+^ antiporter (*mnh*) operon changed dramatically between the methane/sulphate condition (mean ranking: 9.7%, CH_4_) and CO/sulphate/bicarbonate treatments (mean ranking 86.5%, 0.1 bar CO), representing an average 188-fold increase in expression (0.1bar CO vs. CH_4_); (Fig. 7, Extended Data Fig. 8, Supplementary Data 2). Other genes such as the Flx-Hdr complex, a key component in the cycling of DsrC in the CO metabolism in SRB cells (*29*), showed comparable expression levels with CO vs. methane (Fig. 7, Extended Data Fig. 8, Supplementary Data 2). SRB genes previously identified to be involved in syntrophic electron transfer and carbon fixation (via Wood-Ljungdahl pathway) also showed equivalent expression levels in treatments with 0.1 bar CO vs. CH_4_/sulphate (Fig. 7, Extended Data Fig. 8, Supplementary Data 2).

## Discussion

The syntrophic ANME-SRB consortia living in deep-sea sediments have been well-characterized for their ability to oxidize methane coupled with sulphate reduction. Although environmental, genomic, and thermodynamic evidence suggests that marine methanotrophic ANME might use alternative electron donors such as CO, this had not been experimentally tested. Using complementary techniques including long-term microcosm experiments, multi-omics, and single cell analyses, we demonstrate that ANME and their syntrophic SRB partners are capable of actively oxidize CO in seep sediments.

### CO oxidation by ANME-2b/syntrophic SRB

We initially confirmed that CO oxidation was carried out by anaerobic methanotrophic ANME-2b/ SRB consortia rather than by co-occurring methanogens or other carboxydotrophic microbes, within the complex and diverse microbial community present within our sediment microcosms. We briefly summarize the lines of evidence from our geochemical, transcriptomics, and single cell isotopic analyses which support this conclusion.

First, we did not recover known methanogens in our microcosm experiments after four months of incubation by 16S rRNA amplicon sequencing (Extended Data Fig. 2). Transcripts from CO dehydrogenases or *mcr* genes phylogenetically associated with methanogenic archaea were also lacking in our metatranscriptomic analysis, which was overwhelmingly dominated by ANME-2b and Seep-SRB1g. Finally, while related CO-oxidizing methanogens like *Methanosarcina* sp. are known to produce acetate and formate as major products along with methane during growth on CO (*10*), neither metabolite was detected in our microcosm experiments. Collectively, these findings reduce the likelihood that non-ANME-SRB microbes had any meaningful contribution to the CO oxidation occurring in our experiments. Instead, these findings support that anaerobic ANME and SRB consortia, specifically ANME-2b/Seep SRB-1g, are catalysing CO oxidation in our seep sediment microcosm experiments.

The second line of evidence came from direct analysis of the ANME-2b and SeepSRB-1g consortia themselves. In experiments where CO was provided as the sole electron donor, CO oxidation occurred alongside the reduction of both sulphate and CO_2_ (bicarbonate), resulting in the production of sulphide and methane. Despite the lack of methane headspace in these treatments, ANME-SRB consortia were confirmed to be anabolically active by FISH-nanoSIMS, while GC-MS analysis revealed the production of methane linked to CO_2_ reduction. Still, the detection of activity by ANME-SRB on its own does not unequivocally identify them as the primary CO oxidizers, as it remained possible that an as yet unidentified seep sediment microorganism could have oxidized CO and generated methane, subsequently fuelling sulphate-coupled AOM by ANME-SRB (e.g., Hadarchaeota MAG, Guaymas_P_008, as previously described in the Guaymas Basin) (*18*). To directly test whether ANME-2b was capable of CO oxidation independent of sulphate-coupled AOM, we incubated the same seep sediments in a sulphate-free medium and evaluated the activity of ANME. Under these conditions with CO_2_ as the sole electron acceptor, we again observed CO oxidation and methane production and, importantly, the ANME-2b cells were anabolically active, supporting their role as the primary CO oxidizer. While previous studies have reported that ANME archaea are not able to oxidize methane with CO_2_ as the electron acceptor (*41*), our results demonstrate that ANME-2b can oxidize CO coupled with methane production, and indicate that ANME-SRB consortia remain active even in the absence of an external methane source.

Physiologically, both catabolic and anabolic activity of ANME-SRB decreased with increasing CO concentration, although cell-specific activity remained high at lower CO concentrations (0.1 bar). This trend is consistent with previous reports of microbial inhibition in response to CO exposure (*10*, *42*) and is also supported by the higher expression levels of stress-related genes under CO conditions. A CO partial pressure of 0.1 bar in the headspace corresponds to 112.83 µM dissolved in seawater at 4°C (*43*, *44*), which is two to three orders of magnitude higher than CO concentrations reported from deep ocean sediments (hundreds of nM) based on data from the International Ocean Discovery Program (IODP) Expedition 385 in the Guaymas Basin (*19*) and from marine coastal sediments (98.3 - 333.7 nM) of the East China Sea (*45*). Thus, it is likely that CO metabolism by anaerobic methanotrophs may be more pronounced in the natural environments where CO supply is lower.

### Growth or maintenance?

Methane production by ANME-2b was observed at a rate of approximately one-nineth of the AOM rate in parallel incubations with 0.4 bar CO. This observation raised the question of whether ANME-2b archaea are obligate methanotrophs and if they can sustain anabolism through CO oxidation coupled to methane generation. Here we calculated that the ANME-2b methane production from CO (0.4 bar) and CO_2_ resulted in minimal biomass yield per unit of potential energy gained, around 20 times lower than that of canonical sulphate-coupled AOM. This implies that CO metabolism primarily supports maintenance of ANME-2b activity, rather than enabling significant growth. Several factors may explain the low efficiency of energy utilization for biosynthesis.

Methanogenesis may not be a favourable energy conservation pathway for ANME grown on CO. Studies of CO metabolism by *Methanosarcina acetivorans* have shown that acetate and formate production via the acetyl-CoA pathway (involving substrate-level phosphorylation) is the predominant CO-dependent catabolic pathway, rather than methanogenesis (*10*, *30*). In these studies, more than 75% of the CO was converted to acetate, while less than 10% was released as methane during carboxydotrophic growth. However, in our microcosm experiments with ANME-2b, neither acetate nor formate was detected during CO oxidation. Furthermore, phosphotransacetylase (*pta*) and acetate kinase (*ackA*), two critical genes involved in substrate-level phosphorylation in the conversion of acetyl phosphate from acetyl-CoA to acetate, are absent from ANME genomes (*6*). Another route from acetyl-CoA to acetate is via AMP-forming acetyl-CoA synthetase (Acs) and ADP-forming acetate-CoA ligase (Acd). For that, a previous study proposed that marine lineage ANME-2a might perform acetogenesis via ADP-forming acetate-CoA ligase (Acd) (*46*), which however, generates little energy (4.6 kJ/mol acetate) to sustain microbial life. This was also supported by the enzymatic examination of ANME-2a Acs in a following study, revealing that the conversion of acetate, if existing in ANME, could be most likely used for anabolism rather than energy conservation (*47*). Therefore, energy conservation via the acetyl-CoA pathway is unlikely to be occurring through the same mechanism as reported in these related methanogens.

Another possibility is the presence of an uncharacterized cytoplasmic enzyme associated with the methanogenesis pathway that catalyses the methyl-H_4_SPT reaction without contributing to energy conservation. During CO oxidation, we observed a substantial down-regulation of genes from the methanogenesis pathway (Extended Data Fig. 8). In particular, the N^5^-methyl-H_4_SPT:HS-CoM methyltransferase (*mtr*) operon, which catalyses the energy-conserving (sodium-pumping) methyl transfer reaction, exhibited significantly lower expression under CO conditions (Extended Data Fig. 8, Supplementary Data 1). This suggests a diminished role for the Mtr complex compared with other steps in the methanogenic pathway, potentially explaining the limited energy conservation available for biosynthesis.

The observed ANME-2b expression pattern is consistent with reports of a substantial decrease in *mtr* transcripts for *M. acetivorans* when grown on CO compared to methanol as the energy substrate (*48*). Concurrent with the decrease in *mtr*, the expression levels of the *mts* system in *M. acetivorans* (*mtsD*, *mtsF*, and *mtsH*) was significantly up-regulated under carboxydotrophic conditions (*48*). Based on these findings, Ferry *et al.* proposed a potential bypass for Mtr via direct cytoplasmic methyl transfer at the expense of minimal energy conservation (*49*). Mts genes encode a predicted cytoplasmic methyl-tetrahydromethanopterin:CoM (CH_3_-THMPT:HS-CoM) methyltransferase, which was predicted to serve as an alternative pathway when *mtr* expression is down-regulated under CO conditions. Biochemical enzymatic assays using exogenous protein expression demonstrated CH_3_-THMPT:HS-CoM methyltransferase activity (*50*). However, follow up studies using *in vivo* mutation and physiological experiments did not support this hypothesis, concluding that Mtr is central to methanogenic metabolism and cannot be bypassed easily (*51*).

Supporting this interpretation, homologs to *mts* were not detected in the ANME-2b genome, indicating that the proposed bypass mechanism in (*48*, *49*) is unlikely to compensate the down-regulation of *mtr* in ANME-2b. Consequently, the question of why ANME-2b *mtr* expression is low with CO, and whether this step can be bypassed, remains unresolved. Overall, the absence or inhibition of key energy conservation machinery in ANME-2b under CO conditions likely accounts for the low efficiency of energy conservation for anabolism. Instead, the moderate level of methane production from CO_2_ reduction coupled with CO oxidation may serve as a mechanism for CO detoxification and maintenance of metabolic activity under conditions unfavourable for AOM. Similar low assimilation efficiencies have been reported in other microorganisms, such as sulphate-reducing bacteria during respiration of extracellular metal oxides (*52–54*), and methanogenic hydrocarbon-oxidizing archaea metabolizing long-chain alkanes to methane (*55–57*). CO oxidation by ANME now adds to the list of low carbon assimilation and maintenance metabolisms documented among slow-growing anaerobes.

The question of whether methanotrophic ANME lineages possess net methanogenic capabilities has been a topic of debate for the past decade (*58–61*). To date, there is no direct evidence supporting energy-conserving net methanogenesis in ANME lineages under *in situ* conditions, despite possibility holds that some ANME-1 lineages harbour hydrogenases to potentially support hydrogenotrophic methane production (*6*). However, ANME-2 lineages (e.g., ANME-2b) lack hydrogenases and do not appear to be capable of using H_2_, acetate or methyl compounds as an energy source (*6*). Our experiments instead support CO-dependent methane production by ANME-2b coupled with CO_2_ reduction as a mechanism to sustain basic metabolic activity (e.g., maintenance energy), rather than translating it to sufficient anabolic activity to support cell division at high CO levels (e.g., partial pressures of 0.1 bar or 0.4 bar). In contrast, the methylotrophic methanogen *M. acetivorans* grown with 1 bar of CO was shown to be capable of methanogenic growth, with a generation time of 24 hours (*10*, *30*). Given the lack of growth observed by ANME-2b in our experiments, CO-dependent methane production by these archaea should not be considered a form of conventional methanogenesis, but rather a mode of maintenance and cellular redox balancing. While high CO levels appeared to trigger a stress response in the ANME consortia based on metatranscriptomic data, it is possible that ANME growth rates and metabolism may be more favourable at the lower CO concentrations likely existing in nature.

### CO metabolism in anaerobic methanotrophic consortia

#### CO oxidation and energy metabolism in ANME

ANME-2b expressed CO dehydrogenases (i.e., *cooS*) at a comparable level under CO conditions vs. methane treatments (Fig. 7, Extended Data Fig. 8). A similar trend was also reported in the methanogen *M. acetivorans,* where one *cooS* copy was down-regulated in CO-grown versus acetate-metabolizing cells (*48*). However, more direct enzymatic assays have shown that cultures of CO-metabolizing *M. acetivorans* exhibit approximately three-fold higher CO dehydrogenase activity compared to methanol-grown cultures (*10*). This suggests that ANME-2b may also regulate *cooS* activity at the posttranscriptional or enzymatic level rather than through changes in gene expression when respiring CO.

Carbon monoxide oxidation by CO dehydrogenases in ANME-2b is the initial step in electron flow and subsequently proceeds via various intracellular electron carriers like ferredoxin and F_420_. Concomitant with the cycling of reducing power and maintenance of cellular redox balance, we considered two plausible sinks for CO-derived electrons in ANME-2b archaea: i) through cryptic anaerobic oxidation of methane via Direct Interspecies Electron Transfer (DIET) with syntrophic SRB partners, consistent with its main mode energy conservation (*38*), or ii) via the methanogenesis pathway for CO_2_ reduction (Fig. 8).

**Fig. 8.**
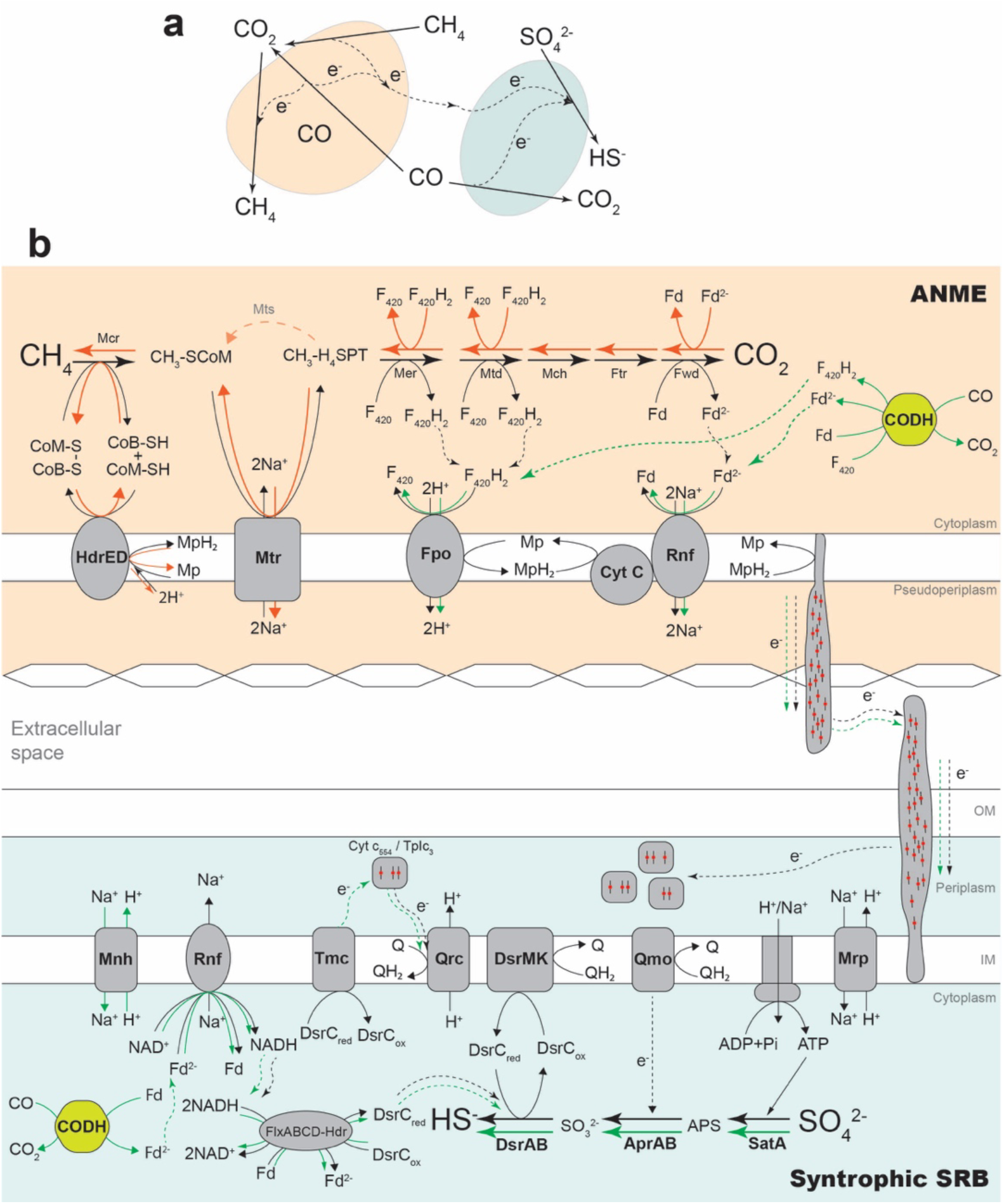
Overview of proposed carbon monoxide oxidation and energy metabolism in ANME-SRB consortia. (**a**) Simple schematic of CO metabolism in ANME-SRB consortia and (**b**) detail of hypothesized energy metabolism in ANME-2b and SRB consortia. In **b**, proposed catalyzing steps connected with black arrows are the conventional energy metabolic pathways of AOM; green arrows indicate the electron flow across ANME-SRB consortia with CO and sulphate as the electron donor and acceptor. CODH, carbon monoxide dehydrogenase (more likely *cooS*); CoM-S-S-CoB, heterodisulfide; F_420_/F_420_H_2_, oxidized/reduced cofactor F_420_; Fd/Fd^2-^, oxidized/reduced ferredoxin; Mcr, methyl-S-CoM reductase; Mtr, N^5^-methyl-H_4_SPT:CoM methyltransferase; Mer, N^5^,N^10^-methylene-H_4_SPT reductase; Mtd, N^5^,N^10^-methylene-H_4_SPT dehydrogenase; Mch, N^5^,N^10^-methenyl-H_4_SPT cyclohydrolase; Ftr, formyl-MF:H_4_SPT formyltransferase; Fmd, formyl-methanofuran dehydrogenase; Hdr, heterodisulfide reductase; HS-CoB, coenzyme B; HS-CoM, coenzyme M; Mp/MpH_2_, oxidized/reduced methanophenazine; Fpo, F_420_H_2_:methanophenazine oxidoreductase; Rnf, rhodobacter nitrogen fixation, energy converting Fd^2-^: acceptor oxidoreductase; Mnh, multisubunit Na^+^/H^+^ antiporters; Tmc, transmembrane channel-like protein; Qrc, quinone-reductase complex; Dsr, dissimilatory sulfite reductase; Qmo, quinol oxidizing complex; Mrp, multiple resistance and pH adaptation antiporters; Flx, flavin oxidoreductase; Apr, adenosine 5’-phosphosulphate reductase; APS, adenylyl-sulphate/adenosine-5’-phosphosulphate; Sat, sulphate adenylyltransferase; Q/QH_2_, oxidized/reduced quinol.

DIET coupled with CO metabolism in ANME-2b is predicted to share the same pathway as methane oxidation (AOM) for shuttling electrons across the cytoplasmic membrane and into the extracellular space (see Fig. 8b, green arrows in the upper panel). In this scenario, membrane-bound enzymes Rnf and/or Fpo reduce ferredoxin (Fd) and F_420_ which subsequently deliver electrons to the membrane-associated electron carrier methanophenazine (Mp). Mp is then predicted to transfer electrons to membrane-bound multi-haem cytochromes *c,* which facilitate electron transfer to the syntrophic partner via DIET (*38*). Notably, both the energy conserving Rnf and Fpo operons were significantly down-regulated under the sulphate-CO condition relative to the sulphate-CH_4_ treatment (Extended Data Fig. 8, Supplementary Data 1), suggesting a metabolic preference for methane. Despite the comparatively low expression levels, however, these pathways appeared to remain active. Therefore, it is conceivable that electrons derived from CO were transferred extracellularly to the syntrophic SRB partner, reducing sulphate as the electron sink for the consortia.

Under the ANME methanogenesis scenario, expression patterns of genes involved in methane production with CO_2_ as the terminal electron sink were examined. Here, electrons carried by electron donors Fd^2-^ and F_420_H_2_ are likely transferred from formyl-methanofuran dehydrogenase (Fwd) to F_420_-dependent N^5^,N^10^-methylene-H_4_MPT dehydrogenase (Mtd) and methylene-H_4_MPT reductase (Mer), following the conventional methanogenesis pathway in *M. acetivorans* (*10*, *30*). Importantly, the ANME-2b *mcr* operon exhibited one of the highest percentile expression rankings in the CO treatments (*ca.* 99.8%, 0.1bar-CO); (Fig. 7, Supplementary Data 1), remaining essentially unchanged relative to the methane condition. The high transcriptional activity of *mcr* is consistent with its involvement in methane generation by ANME-2b and, possibly, in the cryptic oxidation of newly synthesized methane under conditions where both CO and sulphate were provided (Fig. 3).

#### CO oxidation and hypothesized energy metabolism in the syntrophic Seep-SRB1g

Under incubation conditions with sulphate and CO, the syntrophic SRB partner may receive electrons extracellularly with electrons supplied via DIET from CO-oxidizing ANME, and/or intracellularly from direct CO oxidation by SRB via their CO-dehydrogenase. We assess the likely biochemical pathways involved for both possibilities below.

In the first DIET scenario, respiratory electron transfer would follow the same pathways as described for conventional syntrophic sulphate-coupled methane oxidation. Here, electrons are transferred to their syntrophic SRB partner via multi-haem cytochrome *c* proteins, who then couple these electrons to reduce sulphate (*29*). Transcriptomics data from the Seep-SRB1g bacterial partner supports this scenario, with genes encoding extracellular multi-haem cytochrome *c* proteins significantly up-regulated in the syntrophic Seep-SRB1g under CO conditions. This included both *omcX* (fold change: 2.72 (1.0bar-CO *vs.* CH_4_), *P_adj_* = 5.78E-08) and *tmlA* (fold change: 3.15 (1.0bar-CO vs. CH_4_), *P_adj_* = 0.027); (Extended Data Fig. 8, Supplementary Data 2), both predicted to be involved in extracellular electron transfer in SeepSRB-1g (*29*). Electrons delivered from SRB extracellular cytochrome *c* proteins are likely routed through the periplasm and used to reduce quinone pools in the cytoplasmic membrane, and ultimately used for energy generation through dissimilatory sulphate reduction.

As the Seep-SRB1g partner also encodes its own complement of CO dehydrogenases that were actively expressed, it is possible that the syntrophic SRB also directly couple CO oxidation to sulphate reduction (Fig. 7, Supplementary Data 2). CO oxidation in this scenario is predicted to occur through the SRB’s cytoplasmic CO dehydrogenases, serving as the primary source of electrons for downstream sulphate reduction. The direct use of CO would require a physiological shift in the syntrophic Seep-SRB1g partner, where respiratory electron transfer occurs intracellularly rather than, or in addition to, extracellularly via DIET. This change in the direction of electron flow during CO oxidation likely requires metabolic adjustment to optimize the electron transport pathways in the SRB. Here, electrons from CO would be used to reduce ferredoxin, and the reduced ferredoxins, and subsequently NAD^+^, are then used to generate sodium motive force through membrane-bound Rnf and Mnh complexes (*29*). Both Rnf and Mnh were significantly up-regulated by SRB, suggesting these energy conserving complexes were active in treatments with CO and sulphate (Extended Data Fig. 8, Supplementary Data 2). Additionally, Flx-Hdr was also actively expressed at a comparable expression level with CO amendment (Fig. 7, Supplementary Data 2). This complex could serve as a key component in the cycling of DsrC during CO metabolism in SRB, by oxidizing two molecules of NADH to reduce one molecule each of ferredoxin and DsrC (*29*). The predicted coupling by Flx-Hdr effectively recycles ferredoxin from Rnf, while generating reduced DsrC for dissimilatory sulphate reduction. We hypothesize the SRB’s use of ferredoxin, reflected in the high expression of Mnh genes along with the endogenous cycling of DsrC, represents a potential mechanism for the syntrophic Seep-SRB1g to oxidize CO coupled with sulphate reduction independent of electron transfer by ANME (see supplement for additional details). Targeted physiological and biochemical studies of the SRB partner will be necessary for deconvolving their direct and indirect interaction(s) with CO.

The combined evidence from our nanoSIMS experiments and transcriptomics results indicates that syntrophic SeepSRB-1g cells were engaged in both extracellular electron transfer, involving the passage of CO-derived electrons from ANME to SRB, and in the direct oxidation of CO, potentially lessening their energetic reliance on ANME (Table 1, Extended Data Fig. 5, Extended Context). Developing experimental strategies which enable the partial or complete decoupling of metabolic activity by the SRB from its ANME partner, akin to the previously demonstrated use of AQDS to decouple methane-oxidation by ANME from SRB (*8*, *41*), would enable new opportunities in which to investigate the underlying physiology of this enigmatic syntrophic partnership. Future studies using optimized CO concentrations and controlled delivery mechanisms that minimize toxicity may reveal more about the metabolic flexibility of the SRB partners, including the extent to which these syntrophs can respire CO relative to their dependency on ANME.

## Conclusion

In this study, we used microcosm incubations, combined with geochemical analysis, metatranscriptomics, and single-cell activity measurements by FISH–nanoSIMS to demonstrate that anaerobic ANME-SRB consortia can oxidize carbon monoxide (CO) in the absence of methane. Transcriptomic results suggest that CO oxidation in the ANME-2b archaea is catalysed by CO dehydrogenases (likely *cooS*) and used to reduce CO_2_ resulting in methane production through the seven-step methanogenesis/methanotrophy pathway. This redox process appears to be used primarily for maintenance by ANME-2, supporting cellular redox balance, rather than growth. In the presence of sulphate however, the methane generated from CO oxidation in ANME fuels cryptic methane cycling through DIET with their syntrophic sulfate-reducing bacterial partners. Direct CO respiration by the sulphate-reducing bacterial partner also appears likely, allowing for some degree of energetic decoupling between ANME and SRB, while also representing an additional mechanism for energy maintenance within the consortia when methane concentrations may be limiting in the environment. The discovery that environmental AOM consortia can oxidize CO in the absence of methane may enable these consortia to remain viable under changing environmental conditions. This finding deepens our understanding of the complex carbon cycling interactions by these syntrophic archaeal-bacterial associations in deep-sea seep ecosystems.

## Acknowledgements

We thank Stephanie Connon for assistance with 16S rRNA amplicon sequencing; Nathan Dalleska and James Mullahoo for help with GC-TCD, GC-MS, and IC measurements at the Resnick Water and Environment Laboratory (WEL), Caltech; Yunbin Guan for assistance with nanoSIMS operation at the Microanalysis Center for Geochemistry and Cosmochemistry, Caltech; Giada Spigolon in the Biological Imaging Facility in the Caltech Beckman Institute; David Vander Velde for assistance with NMR detection in the Caltech’s Liquid NMR Facility; and Igor A. Antoshechkin and Vijaya Kumar for metatranscriptomic sequencing at the Millard and Muriel Jacobs Genetics and Genomics Laboratory, Caltech. We also acknowledge Emmanuelle Botté for editorial assistance and Dianne Newman and Jared Leadbetter for valuable discussions about this work. This research was supported by the U.S. Department of Energy, Office of Science, Office of Biological and Environmental Research under Award Number DE-SC0022991.

## Disclaimer

This report was prepared as an account of work sponsored by an agency of the United States Government. Neither the United States Government nor any agency thereof, nor any of their employees, makes any warranty, express or implied, or assumes any legal liability or responsibility for the accuracy, completeness, or usefulness of any information, apparatus, product, or process disclosed, or represents that its use would not infringe privately owned rights. Reference herein to any specific commercial product, process, or service by trade name, trademark, manufacturer, or otherwise does not necessarily constitute or imply its endorsement, recommendation, or favouring by the United States Government or any agency thereof. The views and opinions of authors expressed herein do not necessarily state or reflect those of the United States Government or any agency thereof.

## Author contributions

V.J.O. conceived and supervised the project; Y.G. and V.J.O designed the methodology; Y.G. performed the experiments; D.R.U. and Y.G. performed the metatranscriptomic data treatment and analysis; R.M., V.J.O and Y.G. performed the metabolic analysis of ANME-2b and syntrophic SRB; Y.G. generated the original draft of the manuscript in consultation with V.J.O, and all authors provided advice and contributed to the final drafting of the manuscript.

## Materials and methods

### Sediment collection and processing

Methane seep sediments from a white sulphur-oxidizing microbial mat habitat were collected by push core (PC6) with DSV *Alvin* during dive #AD4912 on 27 May 2017 on research cruise AT37-13 on the R/V *Atlantis.* Samples were collected from the Jaco Scar site, off Costa Rica at 1811m water depth (lat 9.1163, lon 284.8372). The sediment core was maintained at 4 °C and processed shipboard upon collection, by upward extrusion and sectioning the sediments into 3-cm-thick horizons which were stored individually in Whirl-Pak bags and then sealed in gas tight mylar bags flushed with argon. These samples were maintained under anoxic conditions at 4 °C until processing back at the shore-based laboratory. Parallel cores were taken for geochemical and microbiological context at this site (*62*).

### Microcosm setup and sampling

All manipulations of the sediment incubations were done anaerobically. Sediments from PC6 (serial number #10079) from Jaco Scar were first divided into two 100 ml serial vials and maintained with sulphate/methane prior to establishment of the formal microcosm experiments. Sediments were evenly allocated into 30 ml serum vials with 2 ml sediment (∼3 g wet weight) distributed per vial and then overlaid with 8 ml artificial seawater to a final volume of 10 ml (1:5 dilution, see Supplemental Materials and Methods). As a killed control, slurried sediment was first fixed with 4% paraformaldehyde overnight and then heat treated at 90 °C for 4 hrs. Electron donors and acceptors were added separately to each incubation bottle. The electron acceptor, Na_2_SO_4_, was added at 5.0 mM final concentration. Methane (1.0 bar, with 5% of ^13^C-methane, Sigma-Aldrich) or ^13^CO (100%, 0.1 bar, 0.4 bar, and 1.0 bar, Sigma-Aldrich) were added as the electron donor with N_2_ as the balance gas to a final total overpressure of 1.0 bar in the headspace. Isotopically labelled ^15^NH_4_^+^ (final concentration 1 mM, 99% ^15^N, Cambridge Isotope Laboratories, Tewksbury, MA, USA) was also added as a tracer for single cell stable isotope probing experiments using FISH-nanoSIMS. All the microcosm incubations were maintained in dark at 4 °C, and 0.5 ml aliquot of the supernatant was taken every 6 days from the bottles using syringes and needles without opening the bottles. All the sampling operations were done in an anaerobic chamber (Coy) to avoid any potential oxidation of compounds of interest in the liquid. 20 µl of the filtered supernatant was immediately added to 380 μl Zn(OAc)_2_ (500 mM) to preserve sulphide as a Zn precipitate for analysis by Cline. Approximately 100 µl of filtered supernatant was dispensed into sealed DIC vials, previously flushed with He and containing 200 µl of H_3_PO_4_ (42.5%). ^13^C-DIC analysis was used to quantify methane consumption in the microcosm experiments (*8*). The remainder of the filtered supernatant was flash frozen in liquid nitrogen and stored at −80 °C for IC analysis (∼200 µl). Microcosm incubations were refreshed with new ASW medium every month prior to sulphate depletion.

Microcosm incubations ran for 4 months. Slurry samples (1 ml from each vial) were taken and fixed in 2% PFA overnight at 4 °C (final concentration 1% PFA). The fixed sediment was subsequently washed 3 times with 3X PBS, followed by a single wash in EtOH/PBS (1:1) and then re-suspended in EtOH/PBS (1:1) to a final volume of 0.5 ml that was used for microscopy and nanoSIMS. The rest of the slurry was flash frozen in liquid nitrogen and kept in −80 °C for DNA extraction.

Microcosm manipulation experiments for monitoring methane production and CO consumption was performed in a separate set of incubations. Here, sediments from the same core (PC6) were mixed with ASW as described in the Supplemental Materials and Methods. Incubations were set up in 10-ml serum vials, under a N_2_ headspace and amended with ^13^CO, and Ar. The addition of Ar gas was used as an internal standard for the GC. DIC samples were taken before GC measurement at the various sampling time points.

### Geochemistry determination

Analysis of sulphide was carried out using the colorimetric methylene blue method in technical triplicate (*63*). Sulphide concentrations were measured using a plate reader (TECAN Sunrise^TM^) by monitoring the absorbance at 670 nm and quantified by comparing to known standards prepared in the same anoxic seawater medium. Quantification of newly formed DIC from ^13^CH_4_ was measured using a GC-IR-MS GasBench II (Thermo Scientific) following protocols described in (*8*). Briefly, the newly formed ^13^C-DIC concentration (Δ[DIC](t_n_)) was calculated from the measured ^13^C fractional abundance (^13^F) following the equation:

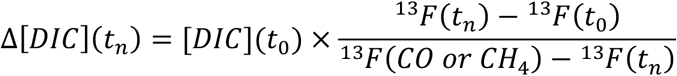

where, [DIC](t_0_) is 5 mM, ^13^F(t_0_) is 0.1153, ^13^F(CO) is 1, and ^13^F(CH_4_) is 0.05. For the total DIC concentration analysis, a standard curve over a series of known concentrations of DIC was determined from 0.1 µmol to 10 µmol. The total DIC concentrations were analysed from the measured peak areas corresponding to the total DIC. For analysis in Figure 4, the newly formed ^13^C-DIC represented the amount of DIC generated from the ^13^CH_4_ oxidation, and the difference between the total DIC and ^13^C-DIC showed the activity of CO oxidation in the incubations. Gas samples were quantified by a GC-TCD equipped with a HP-MOLSIV column (SN US3351826H, Agilent Technologies).

To test whether nitrogenases reduced CO in our microcosm experiments, GC-MS analysis was performed using an Agilent/HP 5890 GC–5972 MSD system equipped with a CP-Molsieve 5A column (Agilent Technologies). Multiple hydrocarbons were monitored including methane (CH_4_), ethylene (C_2_H_4_), ethane (C_2_H_6_) and propane (C_3_H_8_) (hydrocarbon detection limit: low ppb to low ppm range) (*33*, *34*). To test the production of ^12^C- and ^13^C-methane from sulphate-free bottles, a capillary HP-Plot/Q column was used on the same GC-MS system.

NMR spectra were acquired on a Bruker NEO 400 NMR spectrometer. D_2_O was added to aqueous samples to a final concentration of 10% D_2_O. To check for the presence of formate, 1H spectra was acquired using the standard Bruker parameter set WATER, employing a 1D NOESY pulse sequence with a 10 millisecond mixing time. To confirm the identity of formate in our incubations and to estimate its initial concentration, a sample containing formate was spiked with a known formate standard to raise the sample concentration by 100 µM, and the WATER experiment was repeated.

### Extracellular metabolites determination

Extracellular metabolites were quantified via parallel Dionex Integrion HPIC (ThermoFisher, Waltham, MA, USA) ion chromatography systems with anion and cation columns running in parallel housed at the Resnick Water and Environment Lab (WEL) at Caltech. The ion chromatography method was run as previously described (*64*, *65*) with the following modifications.

A Dionex AS-DV autosampler loaded samples diluted 1:50 in 18 MΩ water to serial anion and cation columns and detectors, which are maintained at 30 °C. Anions were resolved by a 2×250 mm Dionex IonPac AS19-4µm analytical column protected by a 2×50 mm Dionex IonPac AG19-4µm guard column (ThermoFisher, Waltham, MA, USA). A potassium hydroxide eluent generator cartridge generated a hydroxide gradient that was pumped at 0.25 mL min^−1^. The gradient was constant at 10 mM for 5 min, increased linearly to 48.5 mM at 27 min, then increased linearly to 50 mM at 40 min. A Dionex AERS 500 suppressor provided suppressed conductivity detection running on recycle mode with an applied current of 31 mA. Cations were resolved by a 2×250 mm Dionex IonPac CS16-4µm analytical column protected by a 2×50 mm Dionex IonPac CG16-4µm guard column. A methanesulfonic acid eluent generator cartridge generated a methanesulfonic acid gradient that was pumped at 0.16 ml min^−1^. The gradient was constant at 10 mM for 5 min, nonlinearly increased to 20 mM at 20 min (Chromeleon curve 7, concave up), and nonlinearly increased to 40 mM at 40 min (Chromeleon curve 1, concave down). A Dionex CERS 2 mm suppressor provided suppressed conductivity detection with an applied voltage of 4.2 V. Chromatographic peaks were integrated by Chromeleon 7.2.9 using the Cobra algorithm and were correlated with concentration by running known standards and generating 4-point calibration curves.

### Sample preparation for aggregate embedding, sectioning, FISH, and nanoSIMS

Sample preparation was carried out following a protocol described previously (*38*). Briefly, paraformaldehyde-fixed consortia in the sediment slurry were detached from the sediment particles via gentle sonication on ice (10s sonication/10s break, 3 times, 4 W) with microtip probe (Branson). AOM aggregates were separated using percoll density centrifugation and concentrated onto a 5 μm filter, covered in molten noble agar (3% <u>in</u> 1×PBS), and embedded in glycol methacrylate (Heraeus Kulzer - Technovit® 8100). Sections of *ca.* 1 μm thickness were cut and stretched on a DI water droplet on a polylysine coated slide with teflon wells (Tekdon Inc) and analysed by fluorescence *in situ* hybridization (FISH) on a Zeiss LSM 980 confocal microscope with 63X objective using general archaeal probe, an ANME-2b specific probe, and a sulphate-reducing bacterial probe. FISH probes and hybridization conditions are provided in the Supplemental Materials and Methods. Images of the FISH-stained consortia were collected, and the location of these consortia were mapped for subsequent nanoSIMS analysis as described below.

### NanoSIMS data acquisition

Isotope enrichment data were collected on a CAMECA nanoSIMS 50L in the Centre for Microanalysis at the California Institute of Technology. Gold-coated samples were pre-sputtered with a 70-pA primary Cs^+^ ion beam with aperture diaphragm D1 = 1 until the ^14^N^12^C^−^ ion counts stabilized. Data were collected using a 2.5-pA beam with D1 = 2 and entrance slit (ES) = 2. Four masses were collected corresponding to the ^12^C^−^, ^13^C^−^, ^14^N^12^C^−^, and ^15^N^12^C^−^ ions, for the determination of ^13^C/^12^C and ^15^N/^14^N ratios, respectively, using a tuning protocol described previously (*8*, *38*, *66*). Acquisitions were collected from different-sized square raster areas (ranging from 10 to 35 µm^2^), with 512 × 512 pixels, and 1 to 2 planes collected per area. Dwell time settings resulted in acquisition times of 30 min to 1 hr per plane depending on the size of the raster.

### NanoSIMS data processing

All data processing and analysis were done in Matlab. NanoSIMS.im data files were initially processed using the Look@NanoSIMS Matlab GUI (*67*) to align planes and export raw data. To align isotopic enrichment data of nanoSIMS with the taxonomic information in the space resolution, the processed nanoSIMS raw ^14^N^12^C^−^ and ^15^N^12^C^−^ images were used for the region of interest (ROI) drawing by hand combining with FISH images as reference. This was performed in the software Fresco (Adobe) on an iPad tablet. Regions of acquisitions that contained ANME-SRB cells were outlined based on the manual cell ROIs, and the elemental information was assigned to each ROI, which generated the single-cell level isotopic enrichment of ^15^F = ^15^N^12^C^−^/(^15^N^12^C^−^ + ^14^N^12^C^−^). All representative nanoSIMS images were illustrated by Limage PV-WAVE (v9.00).

### DNA extraction and 16S rRNA iTag community analysis

DNA was extracted from the sediments stored after 4-month incubation using DNeasy PowerSoil Kit (Qiagen, Valencia, CA, USA) following the manufacturer’s instructions. Illumina iTag 16S rRNA gene sequencing protocol was followed (*41*). Briefly, PCR amplification, barcoding and sequencing of the 16S rRNA hypervariable region (V4-V5) was performed using PCR primers 515f and 906r and amplification conditions already described (*68*). Sequencing data were processed using QIIME v1.8.0 (*69*), clustered at 99% sequencing identity using UCLUST v7.0.1001 (*70*), and the taxonomic identity of the most abundant sequence in each cluster was assigned using a custom SILVA database modified from SILVA Ref NR 99 Database Release 115 (*71*, *72*). iTag community analysis was performed using dada2 and R.

### RNA extraction and metatranscriptomic analysis

To obtain fresh material for RNA analysis, incubations using sediments aliquots from PC6 were developed in parallel following the same protocol as mentioned above (section Microcosm setup). For RNA analysis, electron donors included 1.5 bar methane, 0.1 bar-CO and 1.0 bar-CO. Each condition was run in biological triplicate. Sediment samples were taken on day 6 after the start of the microcosm experiment (∼3 g wet weight), immediately flash frozen in liquid nitrogen, and extracted for total RNA the same day.

RNA was extracted using RNA PowerSoil Total RNA Isolation Kit (Qiagen, Valencia, CA, USA). TURBO DNA-free Kit (ThermoFisher) was used to remove genomic DNA and purified using RNeasy Plus MicroKit (Qiagen). rRNA was removed using Pan-Pro (RNA-Seq) riboPOOL 24 reaction Kit (siTOOLs Biotech). Then, RNA was prepared for sequencing using Superscript IV VILO Master Mix (ThermoFisher/Invitrogen). The library was prepared with the Next® Ultra™ II Directional RNA Library Prep Kit (NEB), and sequencing was conducted on a NextSeq2000 (Illumina) platform in the Millard and Muriel Jacobs Genetics and Genomics Laboratory at the California Institute of Technology, generating 2×150 bp paired-end reads with an average insert length of 300 bp. A depth of 40 M reads per sample was sequenced.

Transcript abundance was quantified using kallisto (*73*) against a single reference database consisting of one ANME-2b MAG (Ga0402030_bin1) and one SRB1g MAG (Ga0402030_bin2) generated previously by a single-aggregate sequencing study (*6*, *29*). These MAGs were the best representative as they originate from an adjacent horizon of the same sediment core and are the best-quality pair of ANME-SRB MAGs (99.4% complete, 1.3% redundant and 98.7% complete, 2.2% redundant, respectively). Counts were read into DESeq2 (*74*) in R for normalization and analysis for differential abundance results presented. Percentile rankings of each gene were also converted from the kallisto TPM data on a per-sample basis. Focal genes were annotated with a combination of KEGG (*75–77*) and Pfam databases (*78*) and manual BLAST (*79*) to known sequences.

### CO dehydrogenase homology searches and phylogeny analysis

A local database of genome sequences from ANME and syntrophic SRB was assembled from two previous studies (*6*, *29*) from our lab, representing all the genomes of ANME-SRB consortia available currently. CO dehydrogenase/acetyl-CoA synthase complex subunit alpha (*cdhA*, MA_RS05310) and anaerobic carbon-monoxide dehydrogenase catalytic subunit (*cooS*, MA_RS06785) from *Methanosarcina acetivorans* C2A were used as template to search for homologs in ANME genomes; anaerobic carbon-monoxide dehydrogenase catalytic subunit (*cooS*, DVU_RS09895) from *Desulfovibrio vulgaris* Hildenborough was used to blast against all syntrophic SRB genomes for homologs. Protein sequences were aligned using MUSCLE (*80*) and then a phylogenetic tree was built using IQ-Tree2 using -m MFP with 1,000 bootstraps (*81*). The tree was visualised using iTOL (*82*).

## Data availability

The raw 16S rRNA amplicon sequencing and metatranscriptomic sequencing reads can be found on the National Centre for Biotechnology Information database under BioProject ID PRJNA1227537. Database IDs for each MAG can be found in Supplementary Data 3.

