## Supplementary Information for "Carbon monoxide oxidation expands the known metabolic capacity in anaerobic methanotrophic consortia"

### Contents of supplementary information

**Supplementary Materials and methods**

**Supplementary Context**

Spatial patterns of cellular activity grown on CO.

Detailed description of CO oxidation and hypothesized energy metabolism in the syntrophic Seep-SRB1g.

**Supplementary Table 1.** Summary of microcosm experiments in this study.

**Supplementary Table 2.** The catabolic activity of ANME-SRB consortia with CO, $CH_4$ and sulphate.

**Extended Data Fig. 1.** The full phylogeny of CO dehydrogenase/acetyl-CoA synthase (CODH/ACS) complex alpha subunit (*cdhA*) and monofunctional anaerobic CO dehydrogenase (*cooS*) in the lineages of ANME and syntrophic SRB.

**Extended Data. Fig. 2.** Community analysis of methane seep sediments using samples after 4-month incubation with CO using partial 16S rRNA gene amplicon sequencing.

**Extended Data Fig. 3.** The schematic of a potential electron sink for nitrogenase.

**Extended Data Fig. 4.** GC-MS measurement of methane production from CO oxidation.

**Extended Data Fig. 5.** Activity relationships between ANME-2b and SeepSRB-1g in AOM consortia.

**Extended Data Fig. 6.** The distribution of gene expression of ANME-2b and syntrophic SeepSRB-1g in different treatments.

**Extended Data Fig. 7.** Volcano plots of gene expression for ANME-2b (upper panel, red) and SeepSRB-1g (lower panel, green).

**Extended Data Fig. 8.** Transcriptomic expression of selected genes involved in the energy metabolism in ANME-2b and syntrophic SeepSRB-1g.

**Supplementary Data 1**. Transcriptomic expression of selected genes from ANME-2b genome.xlsx

**Supplementary Data 2**. Transcriptomic expression of selected genes from SeepSRB-1g genome.xlsx

**Supplementary Data 3**. Summary of IDs for 16S rRNA sequencing and metatranscriptomic sequencing on NCBI database.xlsx

#### Supplementary Materials and methods

##### Medium composition

Artificial seawater was used in the study modified from reference (*1*). The composition of the medium was: NaCl 457 mM, $MgCl_2$ 47 mM, $Na^+$-HEPES (pH 7.5) 25 mM, KCl 7.0 mM, $NaHCO_3$ 5.0 mM, $CaCl_2$ 1.0 mM, $K_2HPO_4$ 1.0 mM, $SeO_3^{2-}$ 0.01 µM, $WO_4^{2-}$ 0.007 µM, 0.1% trace element solution, containing per liter: nitrilotriacetic acid 150 mg, $MnCl_2 \cdot 4H_2O$ 610 mg, $CoCl_2 \cdot 6H_2O$ 420 mg, $ZnCl_2$ 90 mg, $CuCl_2 \cdot 2H_2O$ 7 mg, $AlCl_3$ 6 mg, $H_3BO_3$ 10 mg, $Na_2MoO_4 \cdot 2H_2O$ 20 mg, $SrCl_2 \cdot 6H_2O$ 10 mg, NaBr 10 mg, KI 70 mg, $FeCl_3 \cdot 6H_2O$ 500 mg, $NiCl_2 \cdot 6H_2O$ 25 mg.

##### FISH conditions and probes for this study

The FISH hybridization followed a published protocol (*1*). The samples were hybridized in hybridization buffer containing 35% formamide and incubated at 46 °C for 2 hrs, followed by a wash step at 48 ºC for 15 min.

The FISH probes were used in this study (5 ng/µl for each):

ARCH915 (Alexa488), dual labelled: 5’ to 3’= GTGCTCCCCCGCCAATTCCT (*2*);

ANME-2b-729 (Cy3), dual labelled: 5’ to 3’= CGTTCTCGTAGGGCGCCT (*3*);

DSS658 (Alexa647), dual labelled: 5’ to 3’= TCCACTTCCCTCTCCCAT (*4*, *5*).

#### Supplementary Context

##### Spatial patterns of cellular activity grown on CO

The spatial patterns of activities from ANME-2b and syntrophic SRB were examined (Extended Data Fig. 5). Under CO/sulphate/bicarbonate conditions, ANME-2b cells on average were more active ($^{15}N$ atom % enrichment) than their corresponding SRB partners within each consortium, which differs from the standard methane/sulphate condition where the average activity of archaeal and bacterial populations were correlated at approximately 1:1 as demonstrated previously (*1*, *3*) (Extended Data Fig. 5a). This discrepancy could be attributed to the utilization of bicarbonate in ANME-2b as an additional electron acceptor other than sulphate when respiring CO as an electron source. Besides, ANME-2b and syntrophic SRB living on CO showed distance-independent trends in cellular activity (Extended Data Figs. 4b, c). This is contrast to what is predicted in the case of syntrophic exchange or sharing of a diffusible intermediate (*6*–*8*), that is, a weak syntrophic interaction in the CO-living consortia.

##### Detailed description of CO oxidation and hypothesized energy metabolism in the syntrophic Seep-SRB1g

When CO replaces methane as the electron donor in the system, CO oxidation by CO dehydrogenase in the cytoplasm serves as the primary electron source for both ANME and SRB. Notably, this predicts that respiratory electron transfer in SRB would be stimulated intracellularly rather than extracellularly. The shift in the direction of electron flow likely necessitates metabolic adjustment to optimize electron transport pathways.

CO oxidation is facilitated by CO dehydrogenases, which reduce ferredoxin. Electrons from reduced ferredoxin and $NAD^+$ can also be used in generating sodium motive force through membrane-bound Rnf and Mnh complexes. Mnh functions as a $Na^+/H^+$ antiporter, transporting $Na^+$ or $H^+$ in response to Rnf activity. Notably, *mnh* was significantly upregulated under the CO/sulphate condition, supporting its role in this process. For instance, *mnhB* exhibited a fold change of 19.63 ($P_{adj}$ = 0.00000078) under 0.1bar CO *versus* $CH_4$ condition, while *mnhD* showed the highest increase with a fold change of 593 ($P_{adj}$ = 1.15E-11) (Extended Data Fig. 8, Supplementary Data 2). Percentile ranking profiling revealed little *mnh* expression in the methane condition (~ 0%), but high activity under CO conditions (80-95% of all genes in Seep-SRB1g genome) (Supplementary Data 2). This observation strongly suggests that syntrophic Seep-SRB1g are more dependent on ferredoxin for energy conservation in the presence of CO than methane.

The sodium motive force generated by this process energizes the membrane and is predicted to drive ATP synthesis via ATP synthases as sodium ions are pumped through the membrane. The ATP generated is subsequently used to activate sulphate by Sat, which reduces sulphate to adenosine 5'-phosphosulphate (APS). Electrons from reduced quinones are transferred to AprAB via Qmo complexes, catalyzing the reduction of APS to sulphite ($SO_3^{2-}$). Additional electron transfer occurs through DsrMKJOP complexes, from quinones to DsrC and ultimately to DsrAB, where sulphide ($HS^-$) is generated. The mechanism for DsrC cycling plays an important role in providing an endogenous electron source, as mentioned above.

Flx-Hdr complex serves as a key component in this process, oxidizing two molecules of NADH to reduce one molecule each of ferredoxin and DsrC (*9*). This coupling effectively recycles ferredoxin from Rnf activity, while generating reduced DsrC for dissimilatory sulphate reduction. The preference for ferredoxin, as reflected in the high expression of Mnh genes, along with the endogenous cycling of DsrC, offers a potential mechanism for the syntrophic Seep-SRB1g to live on CO with sulphate.

**Supplementary Table 1. Summary of microcosm experiments in this study.**

| Microcosm | Electron donor | Electron acceptor | Geochemical measurement | nanoSIMS (Y/N) | Result illustration |
|---|---|---|---|---|---|
| Set #1 | $CH_4$ | sulphate + bicarbonate | sulphide, [DIC] | Y | Figs. 2, 6; Extended Data Figs. 2, 5, 6, 7, 8; Tables 1, 2; Supplementary Table 2 |
| | 0.1bar-CO | sulphate + bicarbonate | sulphide, [DIC] | Y | |
| | 0.4bar-CO | sulphate + bicarbonate | sulphide, [DIC] | Y | |
| | 1.0bar-CO | sulphate + bicarbonate | sulphide, [DIC] | Y | |
| Set #2 | 0.1bar-CO | sulphate + bicarbonate | sulphide, [DIC], CO, $CH_4$ | N | Fig. 3 |
| Set #3 | $CH_4$, 0.4bar-CO | sulphate + bicarbonate | [DIC] | N | Fig. 4 |
| Set #4 | 0.4bar-CO | bicarbonate | [DIC], CO, $CH_4$ | Y | Figs. 5, 6; Extended Data Fig. 4; Table 2 |

**Supplementary Table 2. The catabolic activity of ANME-SRB consortia with CO, $CH_4$ and sulphate.**

| ($\mu M\ d^{-1}\ cm^{-3}_{sed}$) | 0.1bar-CO | 0.4bar-CO | 1.0bar-CO | $CH_4$ |
|---|---|---|---|---|
| Sulphide production | 1.33 ± 0.32 | 1.01 ± 0.13 | 0.88 ± 0.23 | 67.63 ± 2.76 |
| Newly formed DIC | 34.97 ± 0.68 | 23.21 ± 4.67 | 12.00 ± 0.07 | 64.63 ± 2.64 |
| Ratio of [DIC production] to [Sulphide production] | 27.48 ± 7.79 | 23.02 ± 3.86 | 14.19 ± 3.60 | 0.96 ± 0.07 |
| Predicted DIC formation from sulphide production | 5.33 ± 1.30[‡] | 4.04 ± 0.53[‡] | 3.53 ± 0.90[‡] | 67.63 ± 2.76[€] |

[‡] The calculation was based on the theoretical redox reaction from Eq. 1.

[€] The calculation was based on the reaction: $CH_4 + SO_4^{2-} = HCO_3^- + HS^- + H_2O$, $\Delta G = -17$ kJ/mol $CH_4$.

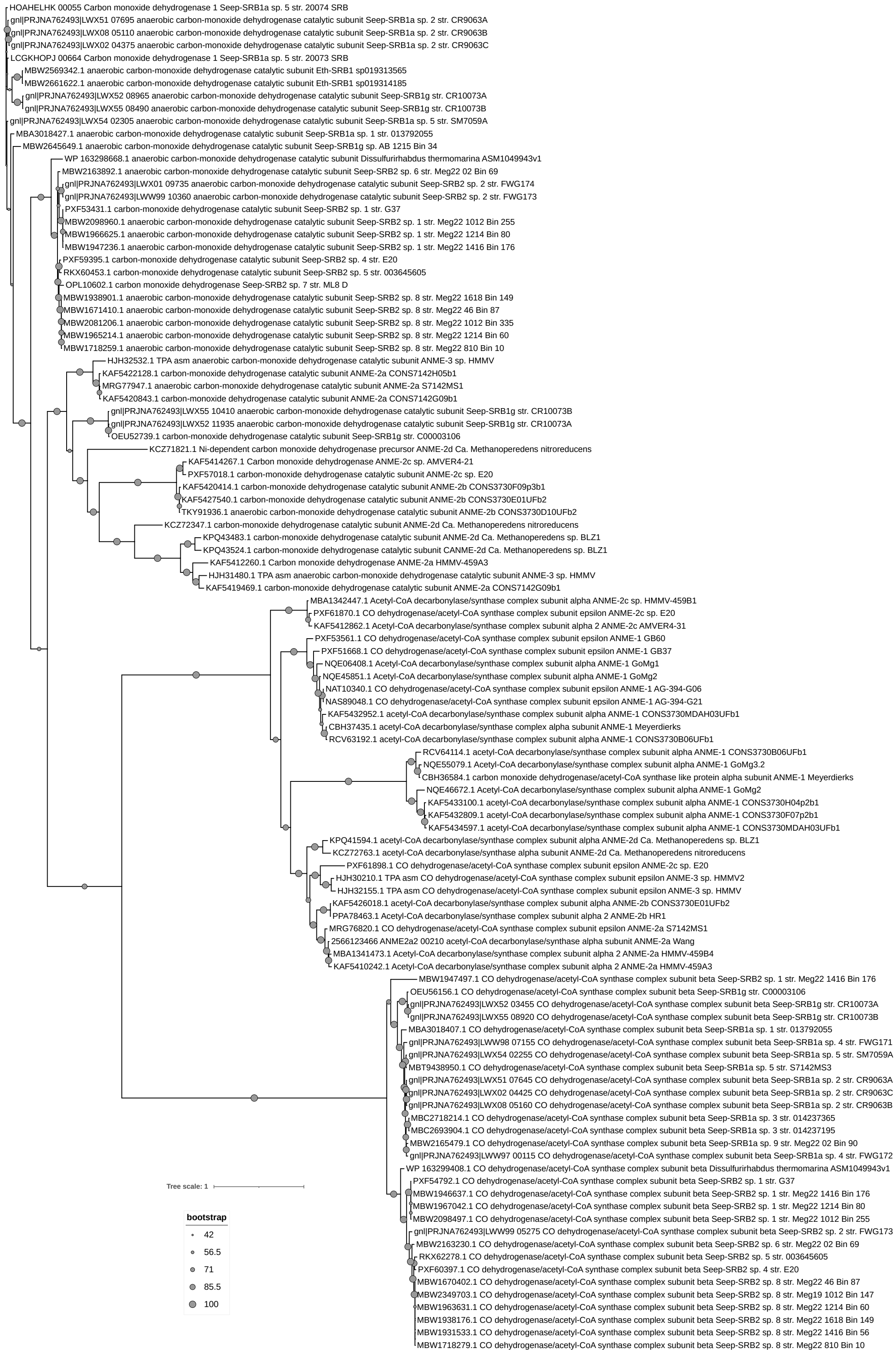


**Extended Data Fig. 1. The full phylogeny of CO dehydrogenase/acetyl-CoA synthase (CODH/ACS) complex alpha subunit (*cdhA*) and monofunctional anaerobic CO dehydrogenase (*cooS*) in the lineages of ANME and syntrophic SRB.**

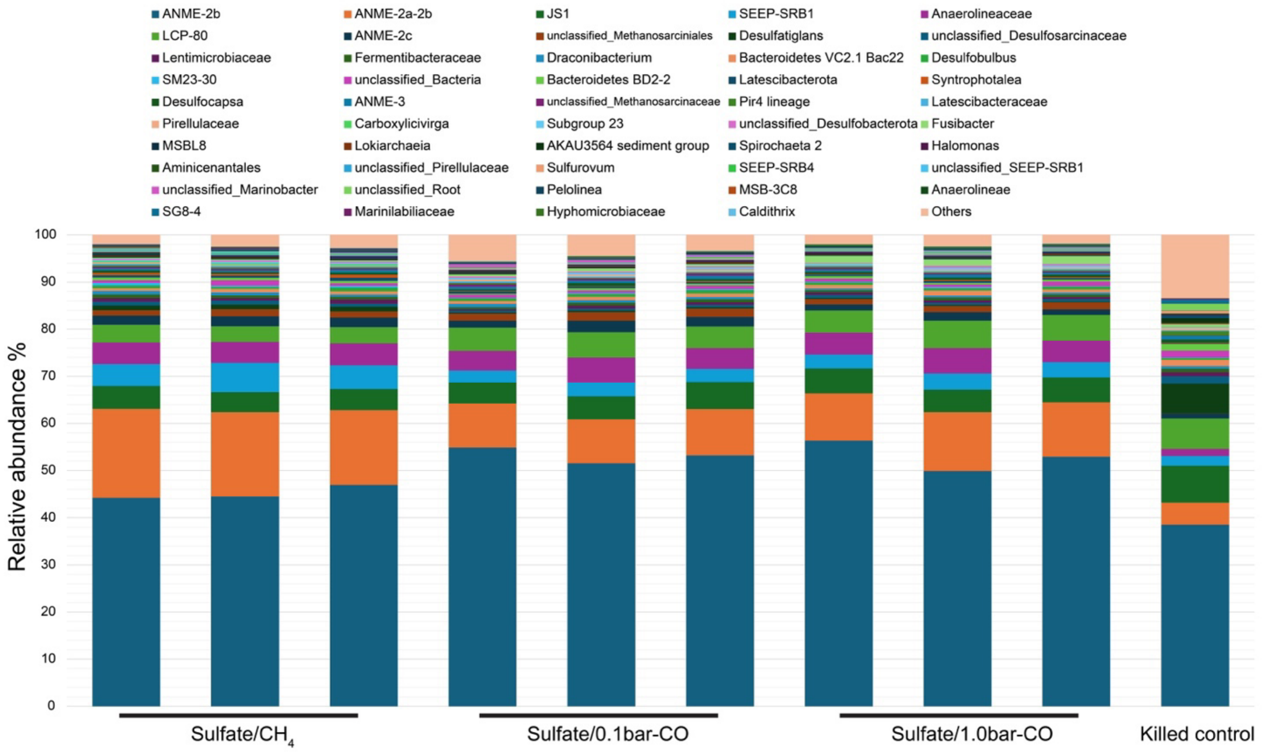


**Extended Data. Fig. 2. Community analysis of methane seep sediments using samples after 4-month incubation with CO using partial 16S rRNA gene amplicon sequencing.** Taxonomic classification is based on the SILVA small subunit rRNA database v138. Taxonomies with more than 1% were shown.

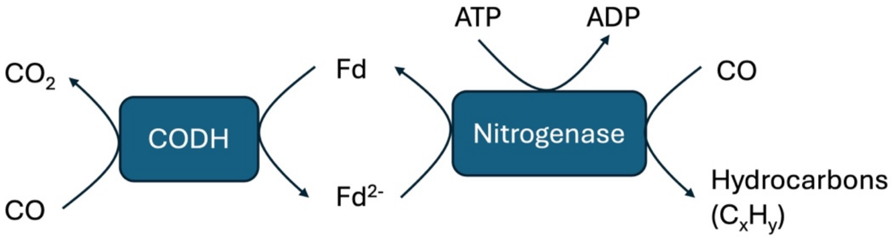


**Extended Data Fig. 3. The schematic of a potential electron sink for nitrogenase.** Electron flow is predicted to start from CO oxidation in CO dehydrogenase, be mediated via electron carrier ferredoxin (Fd), and ultimately feed the CO reducing reaction in nitrogenases with an investment of ATP and result in a series of hydrocarbons as products.

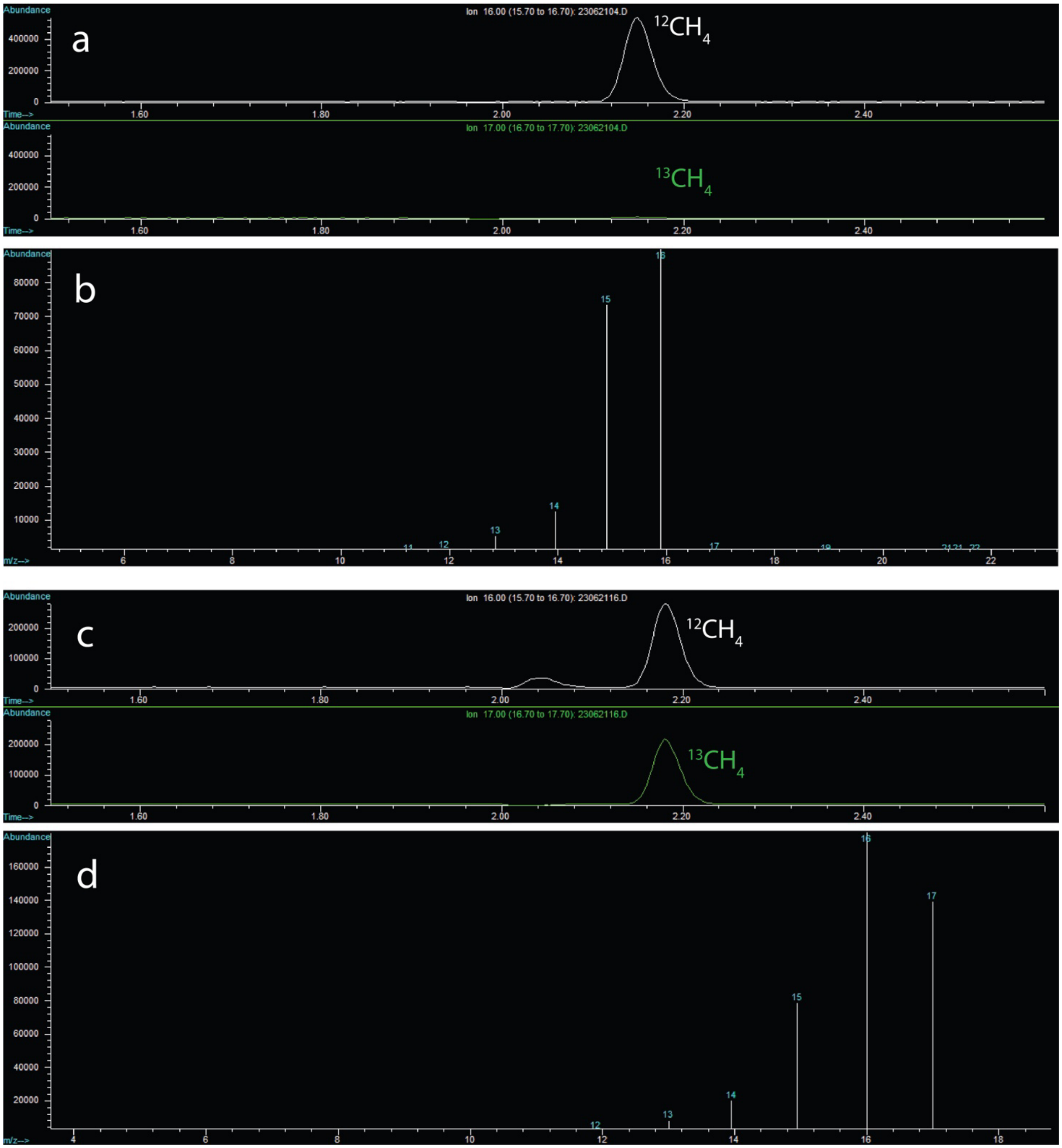


**Extended Data Fig. 4. GC-MS measurement of methane production from CO oxidation. a,** Gas chromatography of $^{12}C$-methane. **b,** The mass-to-charge (m/z) ratios at which $^{12}C$-methane was traced. **c,** A representative gas chromatography of $^{12}C$- and $^{13}C$-methane from CO/bicarbonate maninuplated experiments. **d,** The mass-to-charge (m/z) ratios at which $^{12}C$- and $^{13}C$-methane was traced. In panel **d**, m/z of 17 represents $^{13}CH_4$, 16 for a combination of $^{12}CH_4$ + $^{13}CH_3$, 15 for a combination of $^{12}CH_3$ + $^{13}CH_2$, etc.

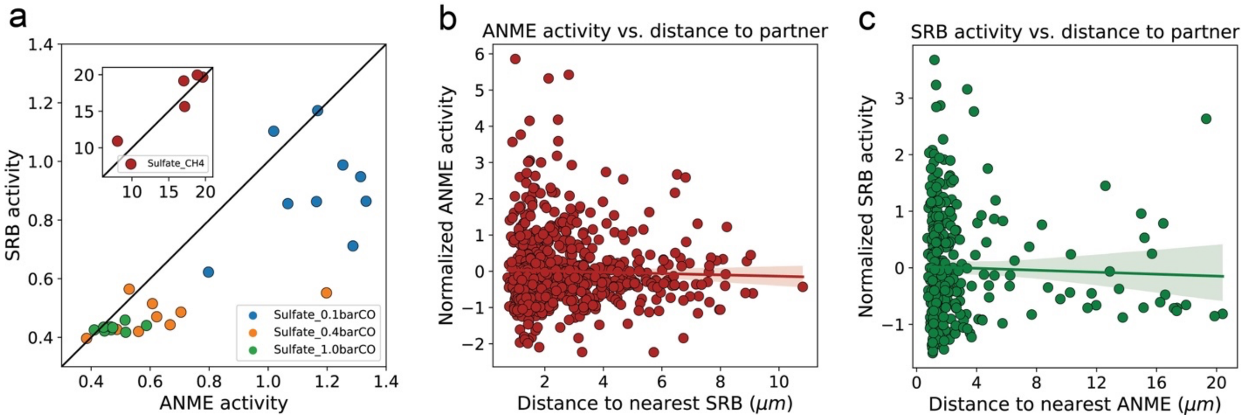


**Extended Data Fig. 5. Activity relationships between ANME-2b and SeepSRB-1g in AOM consortia. a,** Population-level average of SRB activity versus ANME activity for individual AOM consortia. The 1:1 line is shown in black. **b, c,** Normalized activity of single ANME or SRB cells, respectively, plotted against distance to the nearest syntrophic partner. Shaded areas illustrate the 95% confidence intervals in slopes and intercepts of the linear regressions.

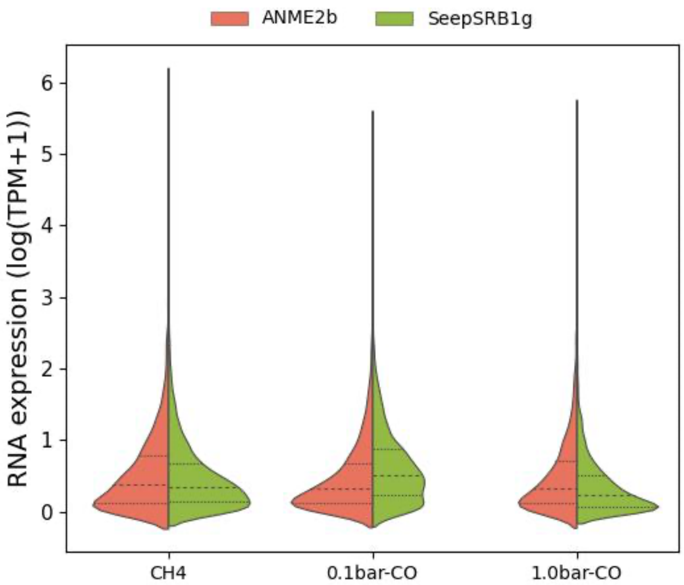


**Extended Data Fig. 6. The distribution of gene expression of ANME-2b and syntrophic SeepSRB-1g in different treatments.** The comparison was based on normalized transcript per million reads (TPM). Dashed lines indicate the quartile of the distribution.

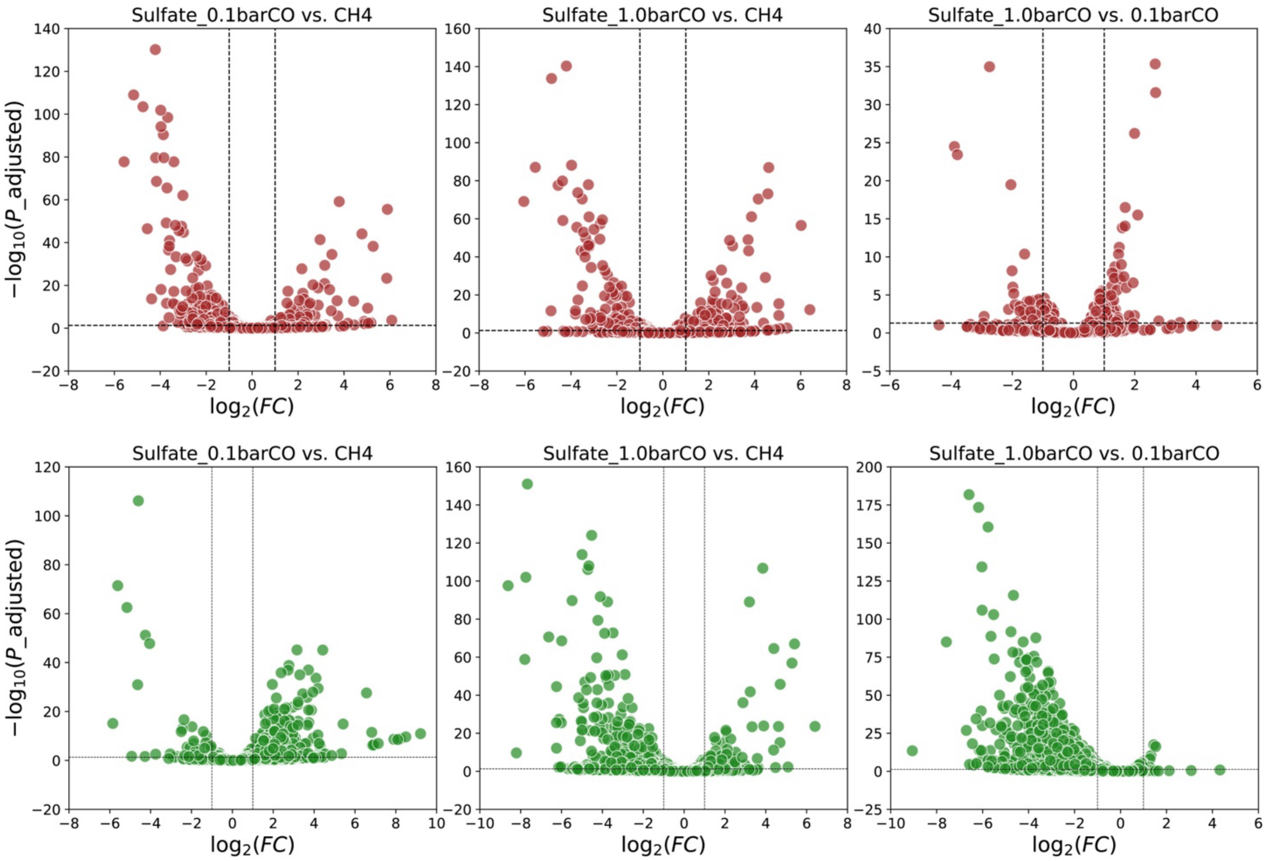


**Extended Data Fig. 7. Volcano plots of gene expression for ANME-2b (upper panel, red) and SeepSRB-1g (lower panel, green).** Conditions include 0.1barCO vs. $CH_4$, 1.0barCO vs. $CH_4$, and 1.0barCO vs. 0.1barCO. The cutoffs were $P_{\_adjusted} = 0.05$ and $\log_2(FC) = 1$.

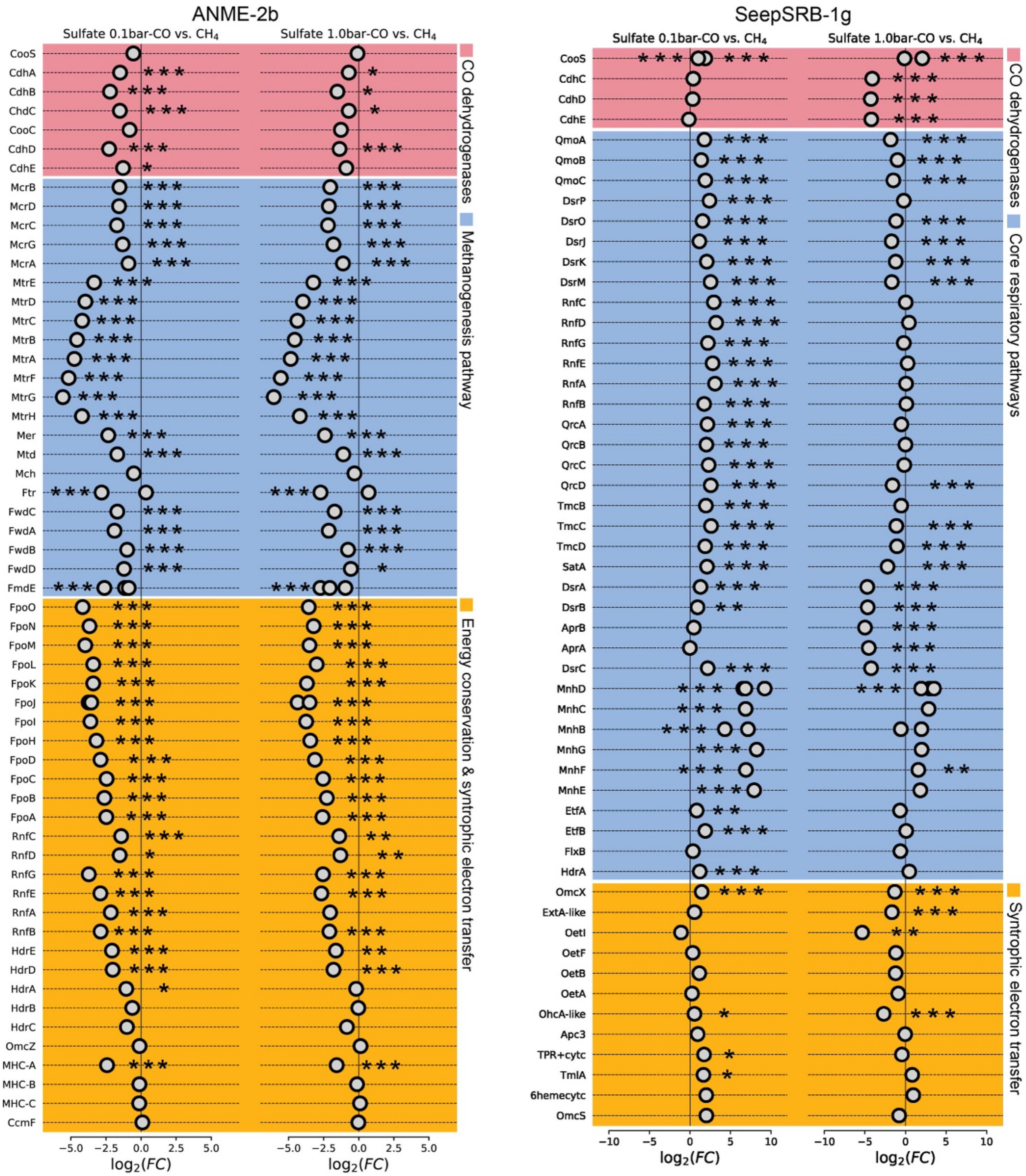


**Extended Data Fig. 8. Transcriptomic expression of selected genes involved in the energy metabolism in ANME-2b and syntrophic SeepSRB-1g.** (**Left**) Selected genes in ANME-2b: CO dehydrogenases (pink), methanogenesis pathway (blue), and energy conservation and syntrophic electron transfer (yellow). (**Right**) Selected genes in SeepSRB-1g: CO dehydrogenases (pink), core respiratory pathways (blue), and syntrophic electron transfer (yellow). The gene expression is centered log-ratio transformed. Differential expression of genes between CO condition and $CH_4$ condition is reported as $\log_2$-fold-change ($\log_2$FC). The pairwise statistical test is based on DESeq2 with the Wald test and adjusted by applying FDR. *$P_{adjusted} < 0.05$; ** $P_{adjusted} < 0.01$; *** $P_{adjusted} < 0.001$. The detailed profiling of those genes is summarized in Supplementary Data 1 and 2.
